# Rice plants alter their yield component parameters based on their tolerance to low temperatures determined by a non-invasive method during the vegetative stage

**DOI:** 10.1101/2023.02.08.527700

**Authors:** Michelini María Carolina, Ayelén Gazquez, Mariana Leonela Checovich, Santiago Javier Maiale, Ana Bernardina Menéndez, Andrés Alberto Rodríguez

**Affiliations:** Laboratorio de Fisiología y Asistencia al Mejoramiento Vegetal, Instituto Tecnológico de Chascomús (INTECH-CONICET-UNSAM), Escuela de Bio y Nanotecnologías (UNSAM), Chascomús, Argentina; Instituto Cavanilles de Biodiversidad y Biología Evolutiva (ICBiBE), Botánica, Fac. CC. Biológicas, Universitat de València, 46100 Burjassot, Valencia, Spain; Departamento de Biodiversidad y Biología Experimental, FCEyN – UBA, INMIBO-CONICET, Buenos Aires, Argentina

**Keywords:** *Oryza sativa*, cold stress, non-destructive techniques, growth and photosynthetic parameters, panicle weight, percentage of empty grain

## Abstract

The first rice phenological stages are commonly subjected to low temperatures (LT) in temperate crop areas. Here, plants of rice genotypes were analyzed according to their LT tolerance at seedling stage, using a realistic experimental temperatures range, and measuring quantitative and operator-independent parameters, fast-obtainable by non-invasive techniques. The ratios between values of these parameters determined in plants under LT condition with respect to the control condition were statistically analyzed. We have found a discriminant formula based on two parameters that allowed to differentiate between genotypes previously described as tolerant and sensitive to LT, with a very low general classification error. The application of the discriminant formula to score plants under LT, using all genotypes from the Rice Diversity Panel 1 which comprehensively represents the genetic variability of the *O. sativa* species, resulted in LT tolerance levels of its subspecies and subpopulations consistent with previous reports. In turn, a strong association between the LT tolerance score at the seedling stage and the panicle weight in plants under field LT during the vegetative stage, suggested that this scoring could be used as an indirect selection factor of genotypes with good yield traits and suited to LT prone environments.

## 1 INTRODUCTION

The farmers from temperate regions sow rice during spring to avoid cold events at the end of the rice cycle, when grain formation takes place (Farrell et al., 2003). This practice implies that rice must develop, from germination until advanced vegetative stages, under a low temperature (LT) stress range that starts around 20ºC to 12-13ºC (Yoshida, 1981; Allen and Ort, 2001; Quintero, 2009). The main symptoms indicating that rice plants suffer LT stress are vegetative growth reduction and alteration of the foliar phenology (Afza et al., 1999; Shimono et al., 2002; Fujino et al., 2004; Ali et al., 2006; Farrell et al., 2006; Baruah et al., 2009; Tavakkoli et al., 2012). In addition, it has been shown that LT stress applied to the rice plants at the vegetative stage, negatively affects the spikelet fertility, which strongly determines grain yield (Shimono et al., 2007). Therefore, to overcome potential losses, it is essential to screen and select genotypes with better LT tolerance to be used as parents in breeding programs. Genotypes with improved LT tolerance obtained would increase yields in an environmental and economically sustainable manner. However, for selective breeding to be effective, there must be genetic variation present in the screened population (Hill, 2001). In the case of *O. sativa*, the genetic diversity is encompassed by two subspecies and 5 subpopulations. The subspecies *japonica ssp*. groups 3 subpopulations, temperate japonica, aromatic and tropical japonica and the subspecies *indica ssp*. groups 2 subpopulations, indica and aus (Garris et al., 2005). These 5 subpopulations are well represented in the Rice Diversity Panel 1 (RDP1) germoplasm collection. With the 413 genotypes contained, this panel spans a wide genetic variability based on the diversity of growth environmental conditions at origin and breeding histories of *O. sativa* accessions (Zhao et al., 2011; Eizenga et al., 2014).

Screening for better stress tolerance and higher yields under field conditions often imposes troubles due to the variability of climate, edaphic conditions and management practices (Afza et al., 1999; Tavakkoli et al., 2012). In parallel, environment-controlled culture chambers are often limited in volume and number, depriving the researcher of the possibility to carried out, fast, cheap and efficiently, large scale experiments with plants completing their life cycle. Therefore, the development of physiological indicators of LT tolerance in seedlings grown indoors that correlate with higher levels of yield components in the field, could be a useful indirect approach for mass screening of a population as big as the RDP1 (Ye et al., 2009). The use of indirect selection criteria in breeding for better yields under stress conditions (drought) was reported for wheat (Passioura, 2002), rice (Babu et al., 2003; Shen et al., 2001) and sugar beet (Ober et al., 2005). However, for indirect selection to be successful, the specific traits must have high correlation with yield (Richards et al., 2001). Field experiments have shown that high yield in rice is causal related to the protection of the photosynthetic apparatus, through excess light dissipation (Wang et al., 2014), and to the photosynthetic performance index (Zhang et a., 2015). More recently, additional photosynthesis and growth-related parameters were shown to distinguish between LT sensitive and tolerant rice genotypes, ie.: the net photosynthetic rate (Pn), stomatal conductance (Gs), some OJIP-test parameters and the area below the leaf growth curve (Gazquez et al., 2015; 2018; 2020; Vilas et al., 2020a). The OJIP-test is a technique that depicts in detail the state of structures related to the photosystem II complex (PSII). It includes the maximum photochemical efficiency of PSII (FV/FM), which represents the maximum efficiency with which an absorbed photon results in the reduction of the quinone A, the maximum efficiency of electron transport in PSII beyond reduced quinone A (ΨE0), and the γRC, which may reflect alterations of the active PSII reaction center (RC) through changes in the active chlorophyll associated to the RC (Strasser et al., 2000; Kalaji et al., 2016). Several authors have reported γRC as a sensitive indicator of abiotic stress tolerance, including cold stress, in different plant species (Hu et al., 2016; Arslan et al., 2018; Bordenave et al., 2019).

Canonical discriminant analysis has been used to identify physiological traits that allow differentiating between stress tolerant and sensitive genotypes (Riggs, 1973; Rascio et al., 2012; Abdolshahi et al., 2015; Safari et al., 2018). Results from the CDA may then be used to assign a new genotype or species of unknown tolerance/sensitivity to one category.

Here we performed and easy and non-invasive study on RDP1 plants confronted with LT at vegetative stage, to detect physiological traits allowing to distinguish between LT tolerant and sensitive genotypes, and to gain insight into the association between the level of LT tolerance at the seedling stage and yield component parameters (YCP) at the ripening stage. For this purpose, we used a canonical discriminant approach for searching the combination of the above-mentioned physiological parameters allowing the most effective grouping of a set of rice genotypes already characterized as tolerant or sensitive. The discriminant function was then used to score the genotypes of the whole RDP1 panel, whose LT tolerances or sensitivities are unknown. Finally, we studied the association between genotype score and yield parameters performance.

## 2 MATERIALS AND METHODS

### 2.1 Plant materials

Seeds from a set of 405 RDP1 different genotypes were used, 334 of which represent the 5 rice subpopulations: 56 aus and 82 indica, 93 temperate japonica, 89 tropical japonica, and 14 aromatic genotypes. In addition, the study included 58 admixed and 13 unclassified RDP1 genotypes.

### 2.2 Experiment 1. Characterization of RDP1 genotypes with respect to their tolerance/sensitivity to LT stress at vegetative stage

#### 2.2.1 Plant material and growth conditions

Among the set of RPD1 genotypes, there were 24 and 25 ones, previously characterized respectively as cold tolerant and sensitive by other authors, which therefore were use as reference (Table S1).

Seeds from all genotypes were sown in Petri dishes, on two layers of Whatman Nº 5 filter paper, rinsed with 7 mL carbendazim 0.025 %p/v and incubated at 30 °C in darkness during three days until germination. The resultant seedlings were transplanted to plastic pots containing sterile organic soil extract as substrate and cultivated in a growth chamber Percival E-30B (Percival Scientific, IA, USA). The growth chamber was set with and optimal condition (control) of 24ºC day/night temperatures, 12-h photoperiod, 80% relative humidity and 300 μmol photons m^-2^ s^-1^ photosynthetic active radiation (PAR; Figure S1). When the third leaf of the seedling appeared (stage T3), approximately 12 days after sowing (DAS) for all genotypes, half of the pots within each genotype were transferred to another growth chamber set at a LT condition (stress treatment), which differed from the control treatment in that the average daily temperature was 16ºC (20/13 °C, day/night). In each growth chamber, there were 3 pots per genotype (1 seedling per pot). Thereafter, the following parameters were measured on the third leaf of intact plants:

#### 2.2.2 Growth parameters

The leaf length (from the blade-stem junction to the blade apex) of the third leaf was measured with a ruler from T3 until the final length was reached (Tf; about 1 week to ten days after T3). Leaf length data as a function of time were adjusted to different polynomial functions according to Zhu and Chen (2015), minimizing the overall sum of squared errors (SSE) for each point on the growth curve, using the Solver tool of Excel® software. The lowest SSE was obtained with a polynomial function of degree 5 in all cases. Then, the area value below the growth curve (Area, from here on) was calculated using the Matlab® Software.

#### 2.2.3 Gas exchange parameters

Pn and Gs were measured in the third leaf at Tf using a LED light unit coupled to a portable leaf cuvette (PLC-4) and the photosynthesis system TPS-2 (PPsystems Inc. MA, USA), at light saturation (1500 μmol photons m^-2^ s^-1^ illumination led light).

#### 2.2.4 Chlorophyll a fluorescence emission kinetic and OJIP parameters

The chlorophyll a fluorescence emission kinetic was analyzed in the third leaf at Tf by OJIP-test, 3 h after the photoperiod light stage initiation, using a fluorometer HANDY PEA (Hansatech Instruments® Ltd., King’s Lynn, Norfolk, UK) according to the manufacturer’s instructions. Blade sections of intact leaves were covered with a leaf clip to adapt them to darkness for 20 min. Then, the blade sections were exposed for 3 s pulse of red light (650 nm, 3500 μmol photons m^-2^ s^-1^). The raw fluorescence data of the fluorescence emission kinetic was processed by the PEA plus software (Hansatech Instrument, UK) to determine the different OJIP parameters. The OJIP parameters described above, γRC, FV/FM and ΨE0 were calculated according to the equations described by Bordenave et al. (2019) and Puig et al. (2021).

#### 2.2.5 Data analysis

The ratio between data obtained under the LT condition to that obtained under the control condition (LT/C) was calculated for Area, Pn, Gs, γRC, FV/FM and ΨE0, on the 49 stress reference (sensitive and tolerant) genotypes. Ratios were processed by canonical discriminant analysis (CDA). To obtain the best discrimination between tolerant and sensitive genotypes, several analysis rounds were done, dropping parameters until obtaining the lower classification error. Remaining parameters in the resultant CDA function (F_CDA_) were used to get a LT tolerance score (TS) for each RDP1 genotype, by introducing the corresponding obtained data.

#### 2.2.6 Statistics

Calculated data sets were tested for normality (D’agostino and Pearson, 1973) and subjected to ANOVA and post hoc analyses DGC tests (Di Rienzo et al. 2002) or Student’s t-test (for subpopulations and subspecies, respectively). The Infostat® statistical software package was used throughout the study (Di Rienzo et al., 2018).

### 2.3 Experiment 2. Effect of the LT-induced stress during the vegetative stage, on yield component parameters at the ripening stage

#### 2.3.1 Outdoor temperature

The outdoor mean daily temperature (MDT) of the air was recorded with a Cavadevices® datalogger. The 7-day moving average of MDT (MDT7) was calculated according to Winkler et al. (2018) to smooth the day-to-day fluctuations of the MDT and to estimate the date to relocate the plants in control condition inside the chamber, to a favorable outdoor temperature condition. The MDT7 equation was:

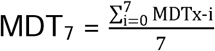

where the letter x represents a particular day after sowing.

#### 2.3.2 Plant material and growth conditions

Sixteen genotypes were randomly taken according to their TS (eight among the 20 most sensitive, and eight among the 20 most tolerant) and used in a second experiment. Seedlings from each of these genotypes were grown until T3 as in Experiment 1 (Figure S1), excepting that 500 (instead of 300) μmol photons m^-2^ s^-1^ PFD from an array with full spectrum COB LED lights were used. Then, half of the pots in each genotype were transferred to the open field (outdoor LT treatment) and half remained within the growth chamber under optimal condition, so that plants were let to grow over the first half of their vegetative stage, either under controlled (optimal) conditions, or under an outdoor LT environment (see registered temperatures in the shadowed range of Figure S2). There were 5 pots (one plant/pot) per each of the 16 genotypes, per treatment. Finally, when the MDT7 exceeded 20ºC during 1 week the upper LT stress condition limit (60 DAS), pots at the growth chamber were transferred to the outdoor (next to LT pots), so that plants from both treatments spent the last part of their vegetative stage and their maturation stage under an environment of favorable temperatures (MDT above the LT stress range, 24ºC average), until they were harvested, and their YCP determined.

#### 2.3.3 Determination of yield component parameters

At harvest, the number of filled grains (NFG), the number of empty grains (NEG), the number of panicles (NP) and the weight of filled grains (WFG) were determined per plant. Then, these measurements were used to calculate the following YCP:

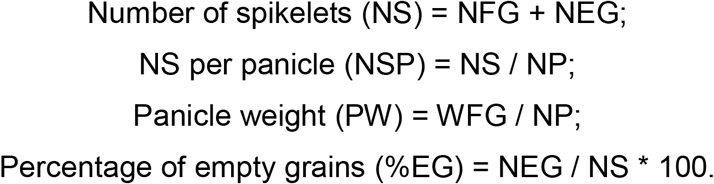

#### 2.3.4 Data analysis and statistics

The LT/C ratio was calculated for all YCP in each plant of the eight genotypes. Pearson’s correlations and linear regressions of these values over their corresponding genotype TS were performed in order to evaluate the association level between LT tolerance levels at the vegetative stage and productivity at ripening. The Infostat® statistical software package was used for this purpose. The most statistically significant linear regression function was used to predict PWLT/C in the remaining RDP1 genotypes.

## 3 RESULTS

### 3.1 Experiment 1. Characterization of RDP1 genotypes with respect to their tolerance/sensitivity to LT stress at vegetative stage

The first step was to determined six parameters related with growth (Area) gas exchange (Pn and Gs) and OJIP-test (γRC, FV/FM and ΨE0) in the third leaf of seedlings from 49 RDP1 genotypes, already characterized as tolerant or sensitive. The LT/C ratio data of the six parameters was analyzed by CDA in order to obtain a function allowing to classify genotypes as tolerant or sensitive, with the lowest error rate. The obtained function (F_CDA_) was defined by the Area_LT/C_ and ΨE0_LT/C_ ratios as follows: F_CDA_ = -7.85 + 4.04 * Area_LT/C_ + 4.02 * ΨE0_LT/C_, with 98.33% of the cases correctly classified (low general classification error = 1.67%, Table S2, Figure 1). The Area_LT/C_ ratio had more influence than the ΨE0_LT/C_ ratio on the discrimination between tolerant (positive TS values) and sensitive (negative TS values) plants, according to their standardized coefficients (1.07 and 0.68, respectively).

**FIGURE 1.**
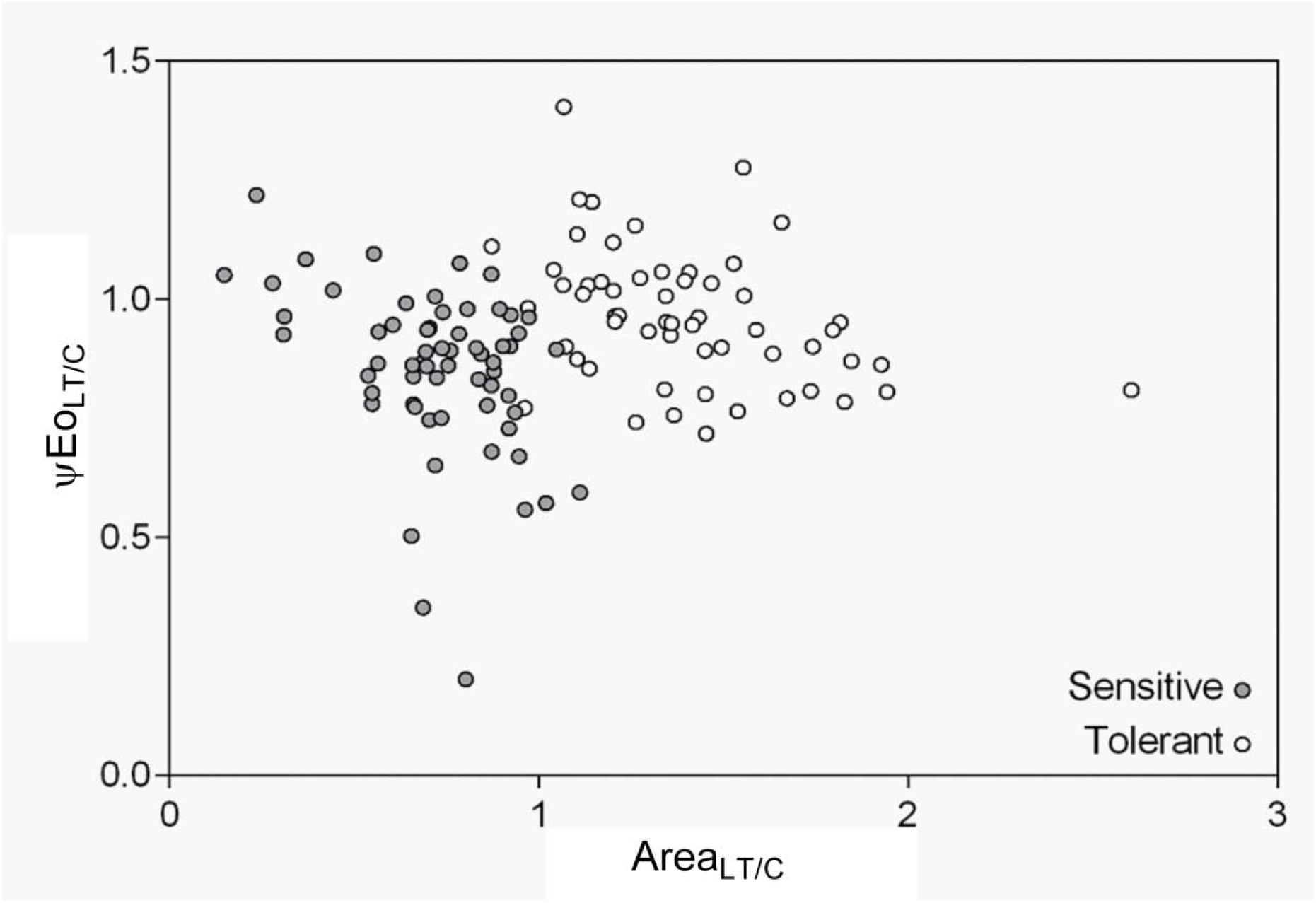
Scatter plot of ΨE0_LT/C_ and Area_LT/C_ ratios from plants belonging to the 49 genotypes previously classified as sensitive or tolerant to cold stress by other authors (see Table S1). Each dot is the average of n = 3 plants per genotype

A TS value for each RDP1 genotype was also calculated by introducing in F_CDA,_ the corresponding Area_LT/C_ and ΨE0_LT/C_ data. Results from scoring all RDP1 genotypes on the basis of their TS showed that overall, around 40% of the genotypes in the panel would be tolerant (positive TS values; Figure 2; Table S3). Within *indica ssp*. and *japonica ssp*. subspecies, tolerant genotypes represented, respectively, 17% and 55% of the total. Analyzing at the subpopulation level, results showed that in average, tropical japonica, aus and indica subpopulations were sensitive, whereas temperate japonica and aromatics were tolerant (Figure 3). In addition, there were not significant differences in the averaged TS within these two subpopulation groups.

**FIGURE 2.**
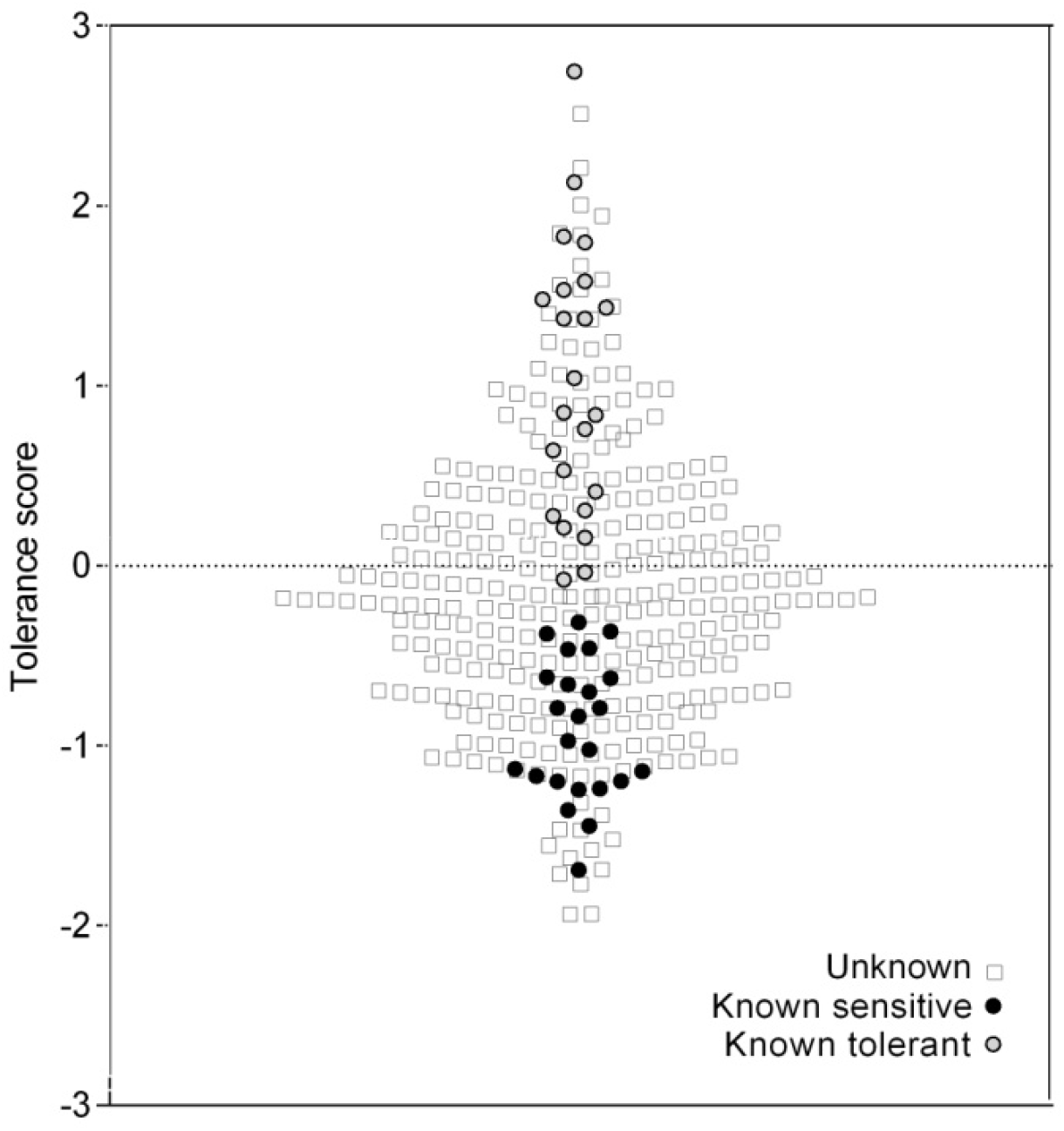
Low temperatures tolerance score for all RDP1 genotypes, calculated from the F_CDA_ function, using the corresponding Area_LT/C_ and ΨE0_LT/C_ data. Grey and black circles are genotypes previously classified as LT tolerant and sensitive, respectively. Empty squares are genotypes whose tolerance/sensitivity was unknown. Each dot is the average of n = 3 plants per genotype

**FIGURE 3.**
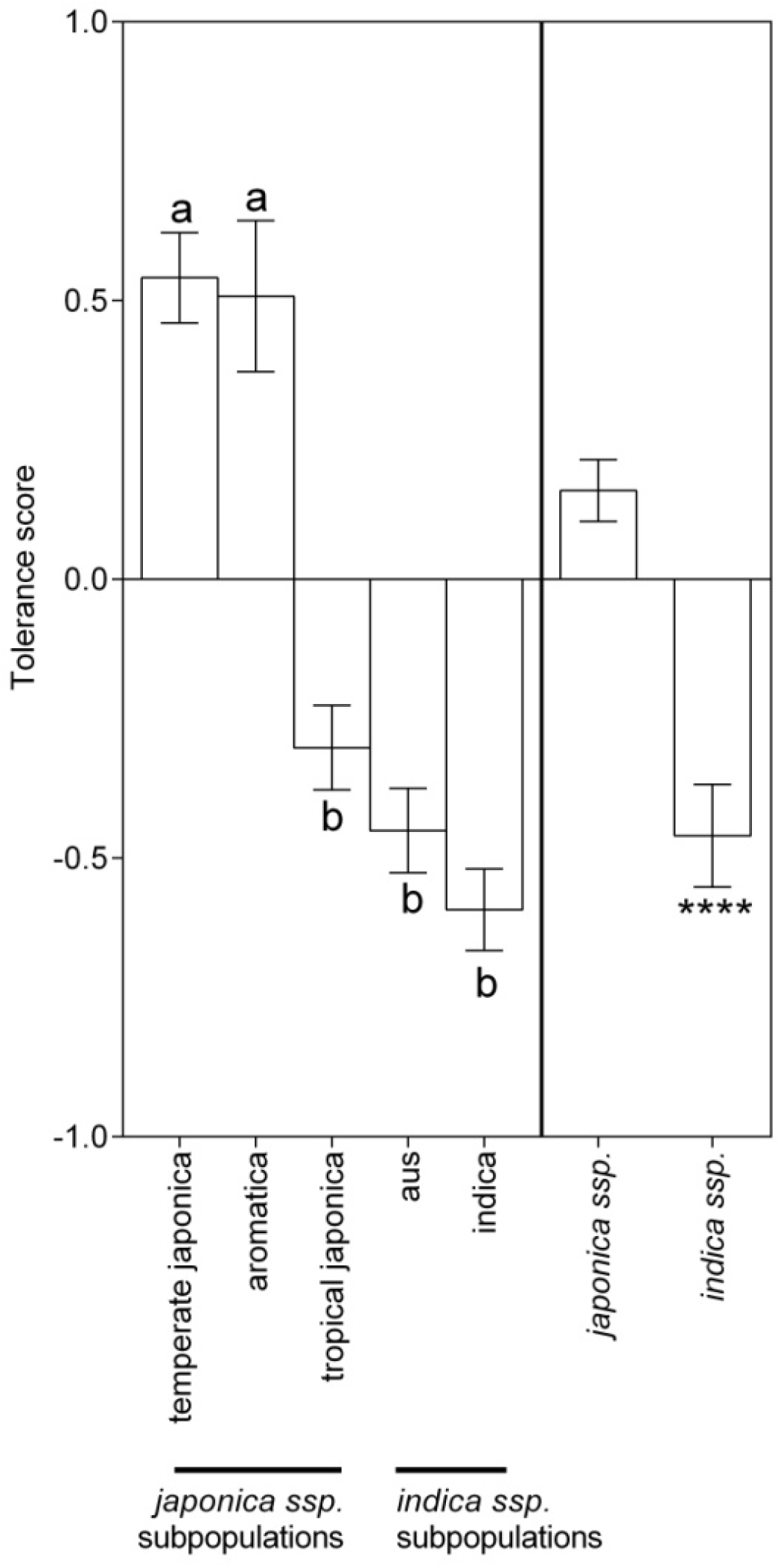
Low temperatures tolerance score in the *O. sativa* population structure. Each bar is the mean ± S.E. of TS for genotypes belonging to the subpopulation. Different letters represent significant differences between subpopulations (ANOVA and post hoc analysis DGC tests *p* < 0.05, n = 3 per genotype; 56, 82, 93, 89 and 14 genotypes, respectively for aus, indica, temperate japonica, tropical japonica and aromatic subpopulations). Asterisks represent significant differences between treatments (Student’s t-test, two samples; \*\*\*\**p* < 0.0001; n = 3 per genotype; 196 and 138 genotypes, respectively for *japonica ssp*. and *indica ssp*.)

### 3.2 Experiment 2. Effect of the LT-induced stress during the vegetative stage, on yield component parameters at the ripening stage

Significantly higher values of PW_LT/C_ and %EG_LT/C_ ratios were found in tolerant, compared with sensitive genotypes (Figure 4A and 4B, respectively), whereas no significant differences due to genotype tolerance/sensitivity were found for the NS_LT/C_ and NSP_LT/C_ ratios (data not shown). In parallel, high correlation coefficients were found between the TS previously determined for the eight randomly taken genotypes with the PW_LT/C_ and %EG_LT/C_ ratios (Table 1). When these last parameters were regressed over the TS, it was found that more than 75% of the PW_LT/C_ ratio variation may be explained by the TS, whereas this relationship was weak for the %EG_LT/C ratio_ (Figures 5A and 5B, respectively). On the basis of the former linear regression function, F = 0.9301 + (0.2612 * TS), it was predicted the PW_LT/C_ ratio (PPW from now on) for all the RDP1 genotypes (Table S4). Grouping the RDP1 genotypes according to their corresponding subspecies revealed that *japonica ssp*. presented in average, an almost 20% higher PPW than *indica ssp*. (Figure 6). When genotypes were grouped by subpopulations, indicated that temperate japonica and aromatica presented similar PPW averages, which were 30% higher than those from tropical japonica, indica and aus. In addition, there were not significant differences in PPW within the last three subpopulations.

**TABLE 1.** Correlation matrix of low temperatures tolerance score with yield component parameters. The matrix correlation presents the Pearson r values and its corresponding *p* values to each correlation between TS, and the ratios %EG_LT/C_ and PW_LT/C_.

|  | TS | %EG<sub>LT/C</sub> | PW<sub>LT/C</sub> |
| --- | --- | --- | --- |
| TS | 1 | 6.60E-04 | 1.40E-05 |
| %EG<sub>LT/C</sub> | -0.76 | 1 | 2.70E-05 |
| PW<sub>LT/C</sub> | 0.87 | -0.85 | 1 |

**FIGURE 4.**
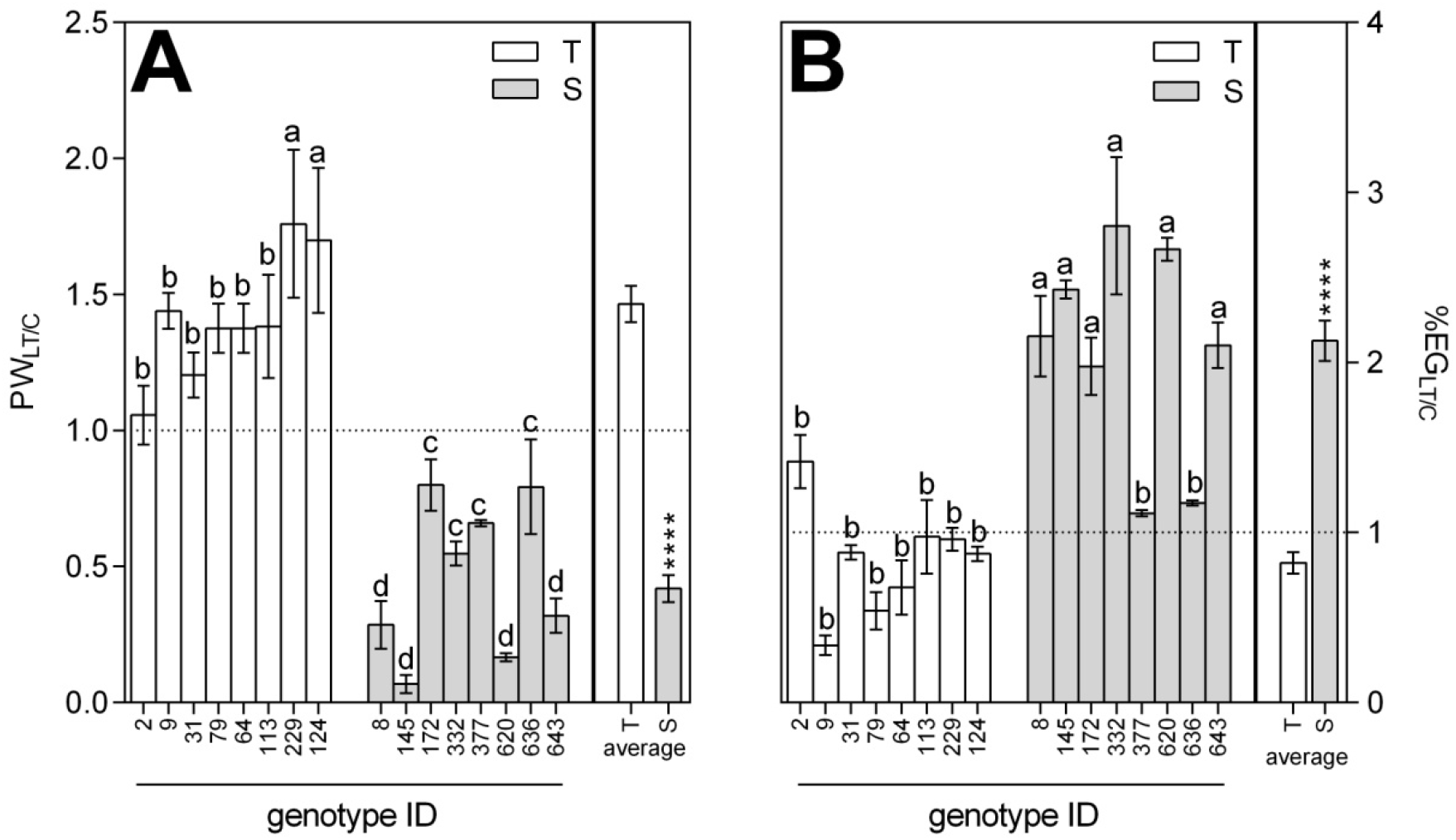
Study of the yield component parameters in sixteen randomly taken genotypes with contrasting low temperatures tolerance score (8/8 tolerants/sensitives). (A) Panicle weight ratio (PW_LT/C_) and (B) Percentage of empty grains ratio (%EG_LT/C_) between LT and control condition. (T) Tolerant (Highest TS values) and (S) sensitive (Lowest TS values) genotypes. Asterisks represent significant differences between treatments (Student’s t-test, two samples; \*\**p* < 0.001; \*\*\*\**p* < 0.0001; data represents mean ± S.E., n = 48; n = 6 per genotype)

**FIGURE 5.**
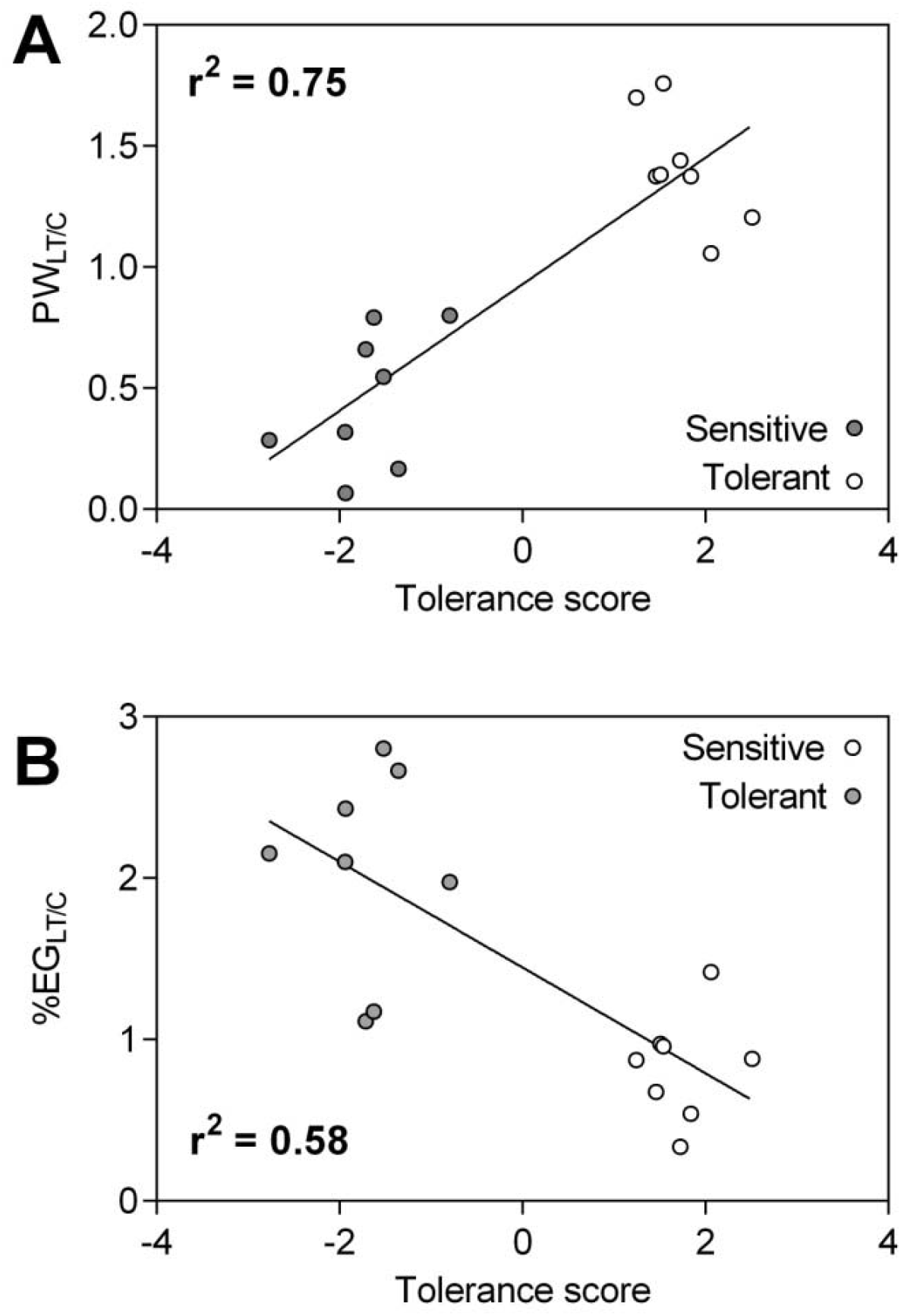
Linear regression analysis of the (A) Panicle weight ratio (PW_LT/C_) and (B) Percentage of empty grains ratio (%EG_LT/C_), between LT and control condition, over the calculated low temperatures tolerance score, on 8 tolerant and 8 sensitive genotypes. The model’s goodness of fit was determined by the coefficient of determination R square (r^2^). The solid line in each graph represents the best regression linear fit model between dependent (Y-axis) and independent (X-axis) variables

**FIGURE 6.**
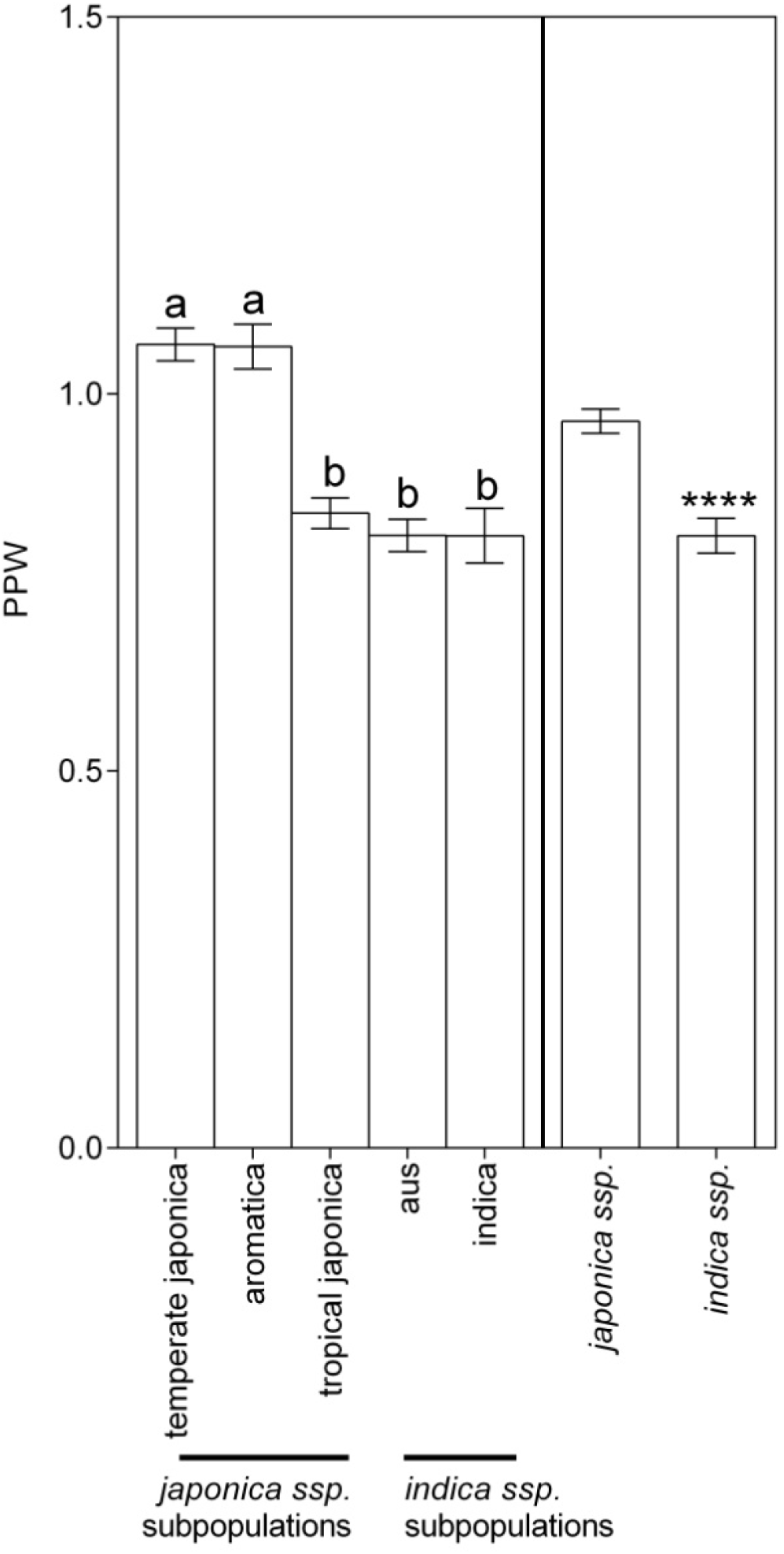
Predicted panicle weight ratio between low temperatures and control conditions in the *O. sativa* population structure. Bars represent means ± S.E. of the PPW for RDP1 genotypes from the 5 subpopulations and 2 subspecies. Different letters represent significant differences between subpopulations (ANOVA and post hoc analysis DGC tests *p* < 0.05, n = 5 per genotype; 56, 82, 93, 89 and 14 genotypes, respectively, for aus, indica, temperate japonica, tropical japonica and aromatic subpopulations). Asterisks represent significant differences between subspecies (Student’s t-test, two samples; \*\*\*\**p* < 0.0001; n = 5 per genotype; 196 and 138 genotypes, respectively, for *japonica ssp*. and *indica ssp*.)

## 4 DISCUSSION

In the present work we followed an approach based on Ye et al. (2009) who suggested that screening for tolerance at the seedling stage could be highly useful for breeding programs directed to improve grain productivity. In doing so, we took into consideration a series of methodological limitations detected in previous works aimed at evaluating LT tolerance in rice.

One trait that differentiates our screening from similar previous ones is the inclusion in the Experiment 1 of almost all accessions of a rice panel (RDP1), which comprehensively represent the *O. sativa* genetic variability. Our results showing an averaged higher LT tolerance in *japonica ssp*., compared with *indica ssp*. confirmed the report by MacKill and Lei (1997), who stated that in general, tolerance and susceptibility to TL in rice are associated with genotypes from *japonica ssp*. and *indica ssp*., respectively. On other hand, our data showing a significantly higher LT tolerance in temperate is in agreement with Garris et al. (2005), who stated that “the two japonica groups represent an adaptive spectrum of an ancient subpopulation from tropical origins to temperate latitudes, with the necessary adaptations to environmental signals such as day length and temperature”.

Our work also confirms previous results suggesting that cold temperatures during the vegetative stage, negatively affect seeds productivity in rice plants (Shimono, 2007; 2011; Matsumura et al., 2012). More importantly, our results showed that the variation in such effect across RDP1 genotypic diversity is a function of the rice plant susceptibility to LT during the first half of its vegetative stage (linear regression models, Figure 5). The last opened the possibility to use this function as a predictive indicator of a final yield for each of the RDP1 genotypes analyzed in Experiment 1 (Table S4).

On other hand, in the 2017/2018 cropping season, Vilas et al. (2020b) carried out a field experiment with RDP1 genotypes, in a location (−32.490254, - 58.348855) within a temperate rice crop area in Argentina. In this location and during that cropping season, the temperatures at the seedling stage and advanced vegetative stages were similar to the LT stress temperatures registered in our Experiment 2 (Figure S3). Plants were cultivated until ripening in a similar way as we did in our study, although there was no control treatment and different YCP were measured, including PW, in 130 RDP1 genotypes. These PW data indicated that PW values were higher in tolerant genotypes than in sensitive genotypes, and in *japonica ssp*. genotypes than in *indica ssp*. genotypes (Figure S4A). These agreed with our results showed in Figures 4A and 6, respectively. But more interestingly, these PW data significantly correlated (*p* < 0.0001) with the TS values that we had assigned to these RPD1 genotypes after Experiment 1 (Figure S4B). Moreover, 4 sensitive and 5 tolerant genotypes from the sixteen genotypes randomly selected for the Experiment 2, retained their tolerance trait in the field study by Vilas and Col., grouping distantly along the correlation line. Last result suggests that our F_CDA_ could be valid as a predictor of PW in plants facing LT during their early vegetative stages.

Although further field studies are necessary to validate our prediction, the information emerging from the present study could be a starting point to select genotypes in breeding programs for the improvement of rice tolerance to LT. In turn, the achievement of genotypes with improved LT tolerance at the vegetative stage would allow to hasten the seeding date. In that way, the grain filling stage, which is the most susceptible one to LT in the rice cycle (Hwang et al., 2019), would coincide with the period of highest light availability and warmer conditions, far from the frequent LT episodes occurring at the end of the rice crop cycle (Quintero, 2009).

Another methodological trait that differentiates our rice germoplasm screening for LT tolerance from previous ones, is the establishment of a control treatment for stress temperature. Actually, the need for studies using systems that provide temperature control had been previously acknowledged by Shimono et al. (2007), who examined the effect of LT before the panicle initiation stage on plant reproduction during panicle development, using the *O. sativa* genotype Sasanishiki.

Most works related to cold tolerance in rice commonly use invasive techniques to obtain data in early phenological stages, what reduces the number of replicates reaching the reproductive stage. In the present work, the Area_LT/C_ and ΨE0_LT/C_ ratios, were the two physiological parameters permiting to differentiate more accurately between genotypes previously described as tolerant and sensitive to LT. Both parameters were measured in seedlings with techniques allowing plants to reach the reproductive stage unharmed, so that we also assessed YCP on them. In addition, measurements could be performed on a high number of plants, in short time. Easy and fast obtainable data is a desirable experimental trait when it comes to breeding programs, as they commonly require analyzing a high number of plants per accession (Galiana-Balaguer et al., 2018).

Another limitation found in previous, similar studies relies on the temperature ranges used for stressing plants. In the vast majority of previous works, plants are stressed using a fixed temperature ranging from 2 to 10ºC (all day long, during several days). These temperatures are below the daily mean critical temperature for rice seedling growth (12-13ºC, according to Yoshida, 1981). Besides, they are rarely registered, or even absent in temperate, cropping rice regions, particularly during the daylight period (Allen and Ort, 2001). To avoid LT that are rather extreme and far from real field conditions, here we used a temperature range that mimics the real day/night LT normally withstanded by rice plants, during the first part of their vegetative stage. This strategy could have helped at preventing intrinsically moderate tolerant genotypes from being classified as sensitive.

It is also common in the literature, the use of stress parameters based on the observer skills such as the leaf color (IRRI 2002; Andaya and Tai 2006; Kim and Tai 2011) the leaf senescence, or the degree of other visual damage signals (Suh et al. 2012) or seed shattering or shedding (Mackill and Lei, 1997). The employement of these type of parameters, along with arbitrary categories (such as low, medium, high, etc) to classify rice plants, opens the possibility of subjective and inaccurate assessments. To avoid this last methodological weakeness, here we utilized the Area_LT/C_ and ΨE0_LT/C_ ratios, two quantitative and operator-independent parameters to score plant genotypes.

All in all, genotypes comprehensively representing the *O. sativa* genetic variability were here scored according to their tolerance/sensitivity to LT stress at an early vegetative stage, using a realistic experimental temperatures range, and measuring a few non-invasive, fast-obtainable and operator-independent parameters.

We have found a discriminant formula based on the Area_LT/C_ and ΨE0_LT/C_ ratios that allowed differentiating between genotypes previously described as tolerant and sensitive to LT, with a very low general classification error (1.67%). The application of the formula to all RDP1 genotypes resulted in LT tolerance levels of the two subspecies and the five subpopulations from the *O. sativa* species compatible with previous reports. In turn, the strong association found between the TS at the vegetative stage and productivity at ripening stage, suggests that this scoring could be used as an indirect selection factor of genotypes with good yield traits and suited to LT prone environments.

## Acknowledgements

This work is part of Michelini M.C. Doctorate. Funding: This work was supported by the Agencia Nacional de Promoción Científica y Tecnológica (PICT Grant Nº 02779).

