## Supplementary material for "Rice plants alter their yield component parameters based on their tolerance to low temperatures determined by a non-invasive method during the vegetative stage": FIGURE S1

**FIGURE S1** Treatments and growing conditions. Characterizations of LT tolerance (Experiment 1) and YCP (Experiment 2). LT: low temperatures; OC: optimal condition

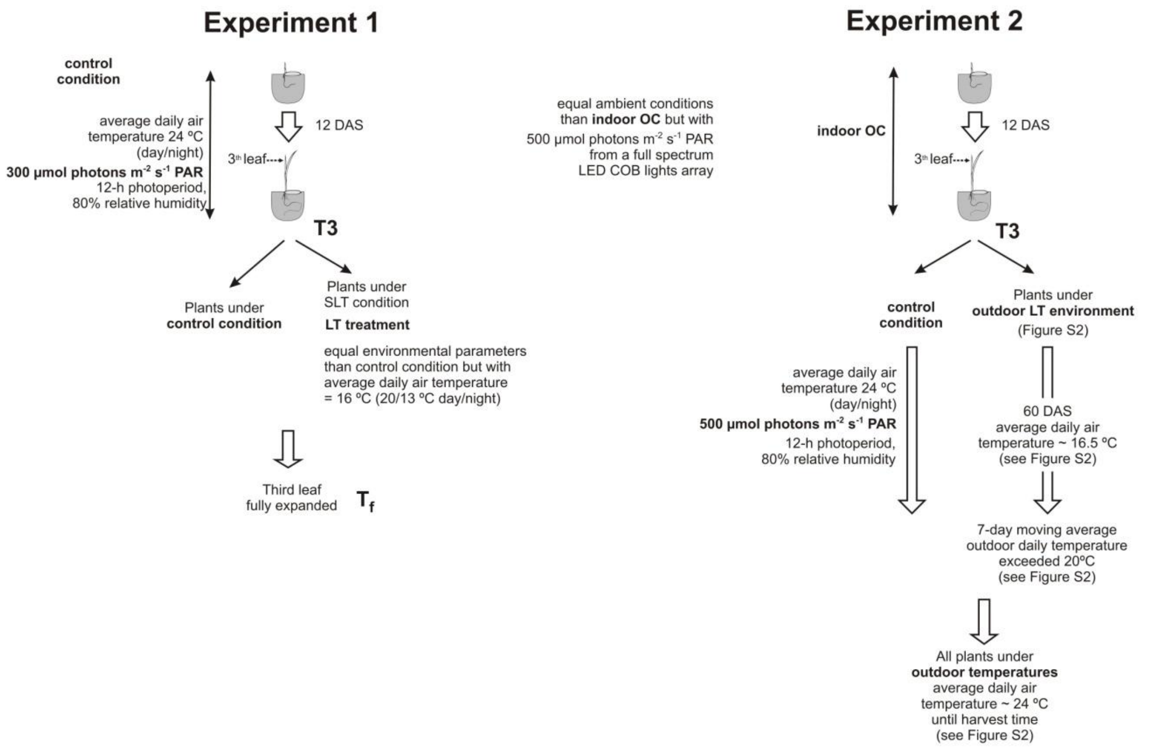
