## Supplementary material for "Rice plants alter their yield component parameters based on their tolerance to low temperatures determined by a non-invasive method during the vegetative stage": FIGURE S2

**FIGURE S2** Mean daily temperature (MDT) and 7-day moving average of the MDT (MDT7) of the air, monitored during the outdoor assay. Arrow 1 (approximately 12 DAS) represents the time when plants growing under optimal condition were transferred to the outdoor LT range during 43 days (Grey area). Arrow 2 represents the time when plants growing under optimal condition were transferred to a favorable outdoor condition, i.e.: when the MDT7 exceeded during 1 week the upper limit (around 20ºC) of the outdoor LT stress condition (after 60 DAS). Arrow 3 represents the date when the first panicle in a certain genotype appeared. Thereafter, all genotypes flowered and produced grains. Harvests proceeded as genotypes produced mature grains, until the last genotype was harvested (130 DAS)

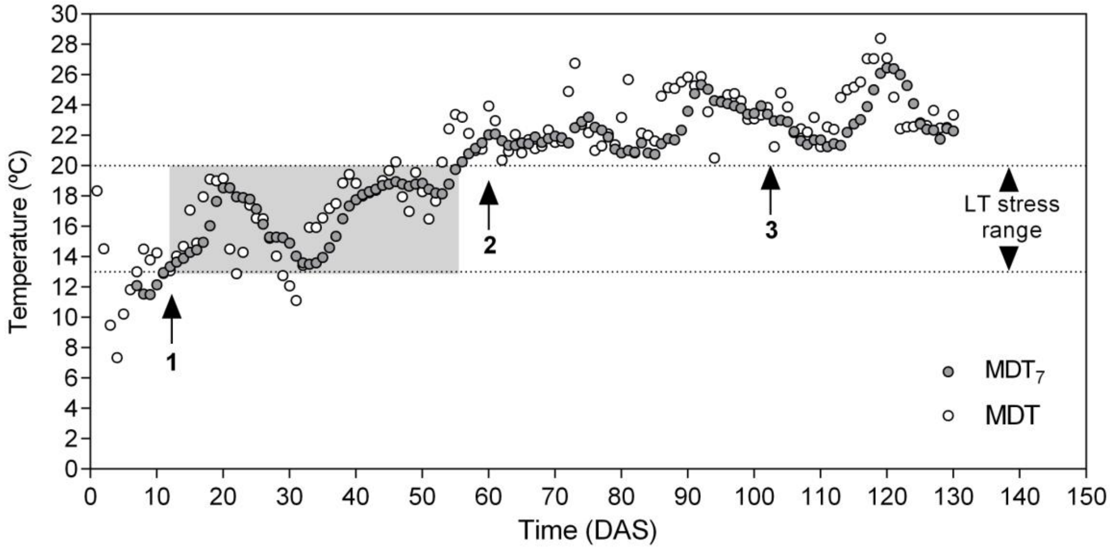
