## Supplementary material for "Rice plants alter their yield component parameters based on their tolerance to low temperatures determined by a non-invasive method during the vegetative stage": FIGURE S3

**FIGURE S3** Mean daily temperature for the assay of Vilas et al. (2020b) during the 2017/18 rice cropping season. The temperature data were extracted from the database of the Agrometeorological Information and Management System belonging to the National Institute of Agricultural Technology. The arrow (approximately 12 DAS) represents the time when plants growing under optimal condition were transferred to the field outdoor conditions

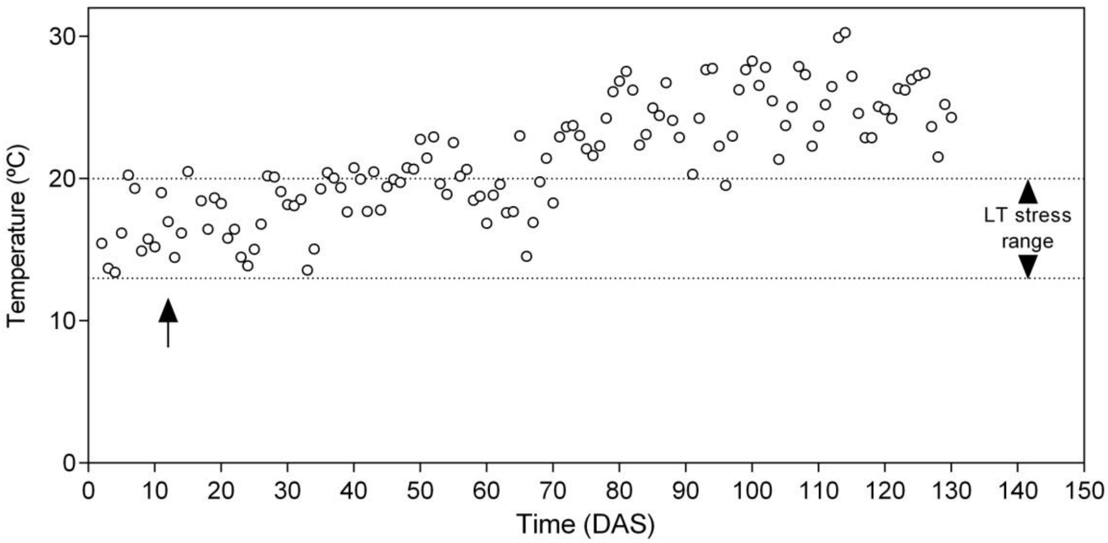
