## Supplementary material for "Rice plants alter their yield component parameters based on their tolerance to low temperatures determined by a non-invasive method during the vegetative stage": FIGURE S4

**FIGURE S4** Panicle weight data from Vilas et al. (2020b). (A) PW data distribution in the *O. sativa* population structure. (B) Association between the data of panicle weight of RDP1 genotypes and their corresponding low temperature tolerance score values from the data of Table S3 in this work. Bars in (A) represent means ± S.E. of PW for RDP1 genotypes from the 5 subpopulations and 2 subspecies. Different letters represent significant differences between subpopulations (ANOVA and post hoc analysis DGC tests $p < 0.05$, n = 3 per genotype). Asterisks represent significant differences between treatments and between subspecies (Student's t-test, two samples; ****$p < 0.0001$; n = 3 per genotype of 130 RDP1 genotypes). The gray and black circles in (B) represent some sensitive and tolerant genotypes, respectively, from the sixteen RDP1 genotypes randomly taken according to their TS represented in Figure 5. The letter r represents the Pearson r from the linear correlation analysis ($p < 0.0001$, n = 130). The solid line represents the best regression linear fit model between dependent (Y-axis) and independent (X-axis) variables

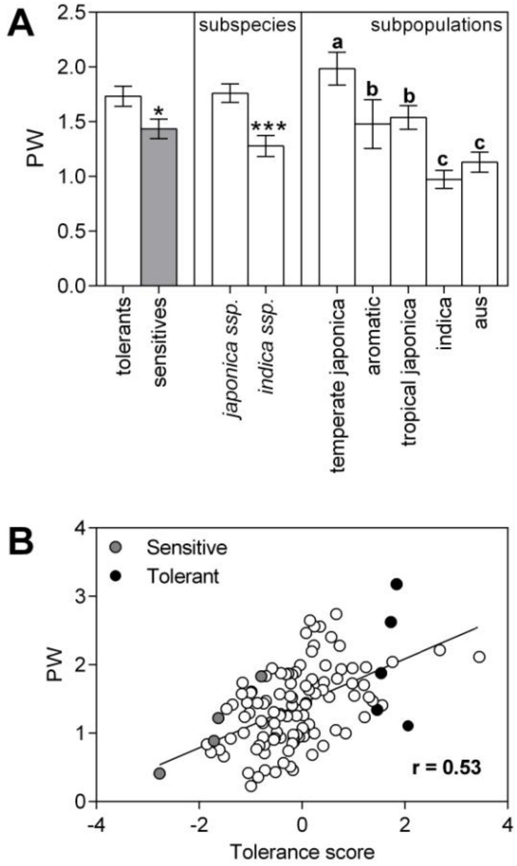
