## Supplementary material for "Rice plants alter their yield component parameters based on their tolerance to low temperatures determined by a non-invasive method during the vegetative stage": TABLE S1

**TABLE S1** RPD1 genotypes previously classified as tolerant and sensitive by different authors.

| RPD1 genotypes name | Subpopulation | Subspecies | cold tolerance classification | Reference |
|---|---|---|---|---|
| Aichi Asahi | temperate-japonica | *japonica ssp.* | tolerant | Mao et al. (2019); Shakiba et al. (2017) |
| Azucena | tropical-japonica | *japonica ssp.* | sensitive | Ye et al. (2009); Shakiba et al. (2017) |
| Baber | temperate-japonica | *japonica ssp.* | tolerant | Mao et al. (2019); Shakiba et al. (2017) |
| Baghlani Nangarhar | temperate-japonica | *japonica ssp.* | tolerant | Mao et al. (2019); Shakiba et al. (2017) |
| Bellardone | temperate-japonica | *japonica ssp.* | tolerant | Shakiba et al. (2017) |
| Beonjo | temperate-japonica | *japonica ssp.* | tolerant | Mao et al. (2019); Shakiba et al. (2017) |
| Binulawan | indica | *indica ssp.* | sensitive | Mao et al. (2019) |
| BJ 1 | aus | *indica ssp.* | sensitive | Mao et al. (2019) |
| Black Gora | aus | *indica ssp.* | sensitive | Mao et al. (2019) |
| Caawa/Fortuna 6-103-15 | tropical-japonica | *japonica ssp.* | tolerant | Mao et al. (2019); Shakiba et al. (2017) |
| Carolina Gold 12033 | tropical-japonica | *japonica ssp.* | tolerant | Shakiba et al. (2017) |
| DZ78 | aus | *indica ssp.* | sensitive | Mao et al. (2019) |
| Erythroceros Hokkaido | temperate-japonica | *japonica ssp.* | tolerant | Goretti et al. (2017) |
| Gogo Lempuk | tropical-japonica | *japonica ssp.* | sensitive | Mao et al. (2019); Shakiba et al. (2017) |
| Gotak Gatik | admixed-japonica | *japonica ssp.* | tolerant | Kim and Tai (2011); Shakiba et al. (2017) |
| Haginomae Mochi | temperate-japonica | *japonica ssp.* | tolerant | Mao et al. (2019); Shakiba et al. (2017) |
| Heukgyeong | temperate-japonica | *japonica ssp.* | tolerant | Mao et al. (2019); Shakiba et al. (2017) |
| IAC 25 | tropical-japonica | *japonica ssp.* | tolerant | Mao et al. (2019); Shakiba et al. (2017) |
| I-Geo-Tze | admixed | *nn* | sensitive | Mao et al. (2019) |
| IRAT 177 | tropical-japonica | *japonica ssp.* | tolerant | Shakiba et al. (2017) |
| Jambu | tropical-japonica | *japonica ssp.* | sensitive | Mao et al. (2019); Shakiba et al. (2017) |
| Jasmine85 | indica | *indica ssp.* | sensitive | Streck et al. (2020) |
| Jaya | indica | *indica ssp.* | sensitive | Ye et al. (2008) |
| JC149 | indica | *indica ssp.* | sensitive | Mao et al. (2019) |
| Kasalath | aus | *indica ssp.* | sensitive | Kim and Tai (2011); Mao et al. (2019) |
| Koshihikari | temperate-japonica | *japonica ssp.* | tolerant | Gazquez et al. (2015), Shakiba et al. (2017) |
| Lemont | tropical-japonica | *japonica ssp.* | tolerant | Cruz et al. (2010) |
| Luk Takhar | temperate-japonica | *japonica ssp.* | tolerant | Mao et al. (2019); Shakiba et al. (2017) |
| M-202 | admixed-japonica | *japonica ssp.* | tolerant | (2017) |
| Ming Hui | indica | *indica ssp.* | sensitive | Mao et al. (2019) |
| MTU9 | indica | *indica ssp.* | sensitive | Mao et al. (2019) |
| N 22 | aus | *indica ssp.* | sensitive | Shakiba et al. (2017) |

| | | | | |
|---|---|---|---|---|
| Nipponbare | temperate-japonica | *japonica ssp.* | tolerant | (2017) |
| Nira | indica | *indica ssp.* | sensitive | Mao et al. (2019) |
| Norin 20 | temperate-japonica | *japonica ssp.* | tolerant | Mao et al. (2019); Shakiba et al. (2017) |
| Oro | temperate-japonica | *japonica ssp.* | tolerant | Shakiba et al. (2017) |
| Ostiglia | admixed-japonica | *japonica ssp.* | tolerant | Shakiba et al. (2017) |
| Padi Kasalle | tropical-japonica | *japonica ssp.* | sensitive | Mao et al. (2019); Shakiba et al. (2017) |
| Pao-Tou-Hung | indica | *indica ssp.* | sensitive | Mao et al. (2019) |
| Pappaku | indica | *indica ssp.* | sensitive | Mao et al. (2019) |
| Phudugey | aus | *indica ssp.* | tolerant | Mao et al. (2019) |
| Pokkali | indica | *indica ssp.* | sensitive | Almeida et al. (2016) |
| RTS4 | indica | *indica ssp.* | sensitive | Mao et al. (2019) |
| Shuang-Chiang | indica | *indica ssp.* | sensitive | Mao et al. (2019) |
| Sintane Diofor | indica | *indica ssp.* | sensitive | Mao et al. (2019) |
| Sultani | tropical-japonica | *japonica ssp.* | tolerant | Wang (2017) |
| Ta Hung Ku | temperate-japonica | *japonica ssp.* | tolerant | Mao et al. (2019); Shakiba et al. (2017) |
| TeQing | indica | *indica ssp.* | sensitive | Mao et al. (2019) |
| ZHE 733 | indica | *indica ssp.* | sensitive | Mao et al. (2019) |
