## Supplementary material for "Rice plants alter their yield component parameters based on their tolerance to low temperatures determined by a non-invasive method during the vegetative stage": TABLE S2

**TABLE S2** Canonical discriminant analysis with the data of $\Psi E0_{LT/C}$ and $Area_{SLT/C}$ ratios in plants belonging to the 49 genotypes previously classified as sensitive or tolerant to low temperatures by other authors according TABLE S1.

| | | | | |
|---|---|---|---|---|
| Variables | | | | |
| $\Psi E0_{LT/C}$ | | | | |
| $AreaS_{LT/C}$ | | | | |
| Classification variable | | | | |
| Low temperatures tolerance score (TS) | | | | |
| Homogeneity test of covariance matrices | | | | |
| Groups | N | Statistic | gl | p-value |
| 2 | 120 | 14,55 | 3 | 0.0022 |
| Eigenvalues | | | | |
| Eigenvalues | % | % accumulated | | |
| 2.51 | 100.00 | 100.00 | | |
| Canonical discriminant functions | | | | |
| Constant | | | -7.85 | |
| $PsiEo_{LT/C}$ | | | 4.02 | |
| $Area_{LT/C}$ | | | 4.02 | |
| Discriminant functions - standardized data with common variances | | | | |
| $PsiEo_{LT/C}$ | | | 0.68 | |
| $Area_{LT/C}$ | | | 1.07 | |
| Cross-classification table (apparent error rate) | | | | |
| Group | sensitive | tolerant | Total | Error (%) |
| Sensitive | 63 | 0 | 63 | 0.00 |
| Tolerant | 2 | 55 | 57 | 3.51 |
| Total | 65 | 55 | 120 | 1.67 |
