## Supplementary material for "Rice plants alter their yield component parameters based on their tolerance to low temperatures determined by a non-invasive method during the vegetative stage": TABLE S3

**TABLE S3** Low temperatures tolerance score for RPD1 genotypes.

| subspecies | subpopulation | NSFTV ID# | tolerance score |
|---|---|---|---|
| *japonica ssp.* | aromatic | 5 | 0.013 |
| *japonica ssp.* | aromatic | 12 | 0.030 |
| *japonica ssp.* | aromatic | 16 | 0.080 |
| *japonica ssp.* | aromatic | 45 | 1.067 |
| *japonica ssp.* | aromatic | 53 | 0.423 |
| *japonica ssp.* | aromatic | 93 | 0.730 |
| *japonica ssp.* | aromatic | 112 | 0.013 |
| *japonica ssp.* | aromatic | 124 | 1.243 |
| *japonica ssp.* | aromatic | 160 | 0.477 |
| *japonica ssp.* | aromatic | 191 | 0.103 |
| *japonica ssp.* | aromatic | 221 | 1.017 |
| *japonica ssp.* | aromatic | 260 | 0.780 |
| *japonica ssp.* | aromatic | 373 | 0.657 |
| *japonica ssp.* | aromatic | 640 | 0.460 |
| *indica ssp.* | aus | 4 | -0.430 |
| *indica ssp.* | aus | 6 | -1.090 |
| *indica ssp.* | aus | 13 | -1.387 |
| *indica ssp.* | aus | 18 | -0.857 |
| *indica ssp.* | aus | 19 | -1.850 |
| *indica ssp.* | aus | 33 | -1.060 |
| *indica ssp.* | aus | 44 | -0.193 |
| *indica ssp.* | aus | 49 | -0.250 |
| *indica ssp.* | aus | 50 | -0.947 |
| *indica ssp.* | aus | 58 | 1.370 |
| *indica ssp.* | aus | 78 | 0.123 |
| *indica ssp.* | aus | 81 | -0.127 |
| *indica ssp.* | aus | 85 | -1.353 |
| *indica ssp.* | aus | 88 | -1.043 |
| *indica ssp.* | aus | 105 | -0.293 |
| *indica ssp.* | aus | 131 | 1.360 |
| *indica ssp.* | aus | 152 | 0.410 |
| *indica ssp.* | aus | 153 | -0.540 |
| *indica ssp.* | aus | 178 | -0.173 |
| *indica ssp.* | aus | 200 | -0.570 |
| *indica ssp.* | aus | 228 | 0.127 |
| *indica ssp.* | aus | 246 | 0.890 |
| *indica ssp.* | aus | 261 | -1.160 |
| *indica ssp.* | aus | 262 | 0.070 |
| *indica ssp.* | aus | 276 | -0.143 |
| *indica ssp.* | aus | 312 | -0.513 |
| *indica ssp.* | aus | 314 | -0.053 |
| *indica ssp.* | aus | 316 | -0.310 |
| *indica ssp.* | aus | 317 | -1.107 |

| | | | |
|---|---|---|---|
| *indica ssp.* | aus | 318 | -0.320 |
| *indica ssp.* | aus | 319 | -0.220 |
| *indica ssp.* | aus | 320 | -0.230 |
| *indica ssp.* | aus | 321 | -1.047 |
| *indica ssp.* | aus | 322 | 0.033 |
| *indica ssp.* | aus | 323 | -1.065 |
| *indica ssp.* | aus | 324 | -0.887 |
| *indica ssp.* | aus | 326 | -0.217 |
| *indica ssp.* | aus | 327 | -0.450 |
| *indica ssp.* | aus | 328 | -0.360 |
| *indica ssp.* | aus | 329 | -0.110 |
| *indica ssp.* | aus | 330 | -0.773 |
| *indica ssp.* | aus | 331 | -0.213 |
| *indica ssp.* | aus | 336 | -1.000 |
| *indica ssp.* | aus | 341 | -0.867 |
| *indica ssp.* | aus | 345 | 0.197 |
| *indica ssp.* | aus | 346 | -0.083 |
| *indica ssp.* | aus | 353 | -0.797 |
| *indica ssp.* | aus | 357 | -0.103 |
| *indica ssp.* | aus | 359 | -0.547 |
| *indica ssp.* | aus | 360 | 0.033 |
| *indica ssp.* | aus | 369 | -0.525 |
| *indica ssp.* | aus | 370 | -0.547 |
| *indica ssp.* | aus | 371 | -0.703 |
| *indica ssp.* | aus | 372 | -0.737 |
| *indica ssp.* | aus | 378 | -0.480 |
| *indica ssp.* | aus | 637 | -1.605 |
| *indica ssp.* | indica | 3 | -0.263 |
| *indica ssp.* | indica | 11 | -0.310 |
| *indica ssp.* | indica | 17 | -1.397 |
| *indica ssp.* | indica | 21 | -0.197 |
| *indica ssp.* | indica | 29 | -0.580 |
| *indica ssp.* | indica | 30 | 0.005 |
| *indica ssp.* | indica | 34 | -0.717 |
| *indica ssp.* | indica | 35 | -0.440 |
| *indica ssp.* | indica | 43 | 0.060 |
| *indica ssp.* | indica | 57 | -0.100 |
| *indica ssp.* | indica | 61 | -0.180 |
| *indica ssp.* | indica | 66 | 0.353 |
| *indica ssp.* | indica | 71 | -0.167 |
| *indica ssp.* | indica | 72 | -0.190 |
| *indica ssp.* | indica | 74 | -1.117 |
| *indica ssp.* | indica | 76 | -0.993 |
| *indica ssp.* | indica | 77 | -1.327 |
| *indica ssp.* | indica | 90 | -0.060 |
| *indica ssp.* | indica | 97 | -0.983 |
| *indica ssp.* | indica | 102 | -0.753 |

| | | | |
|---|---|---|---|
| *indica ssp.* | indica | 106 | -1.517 |
| *indica ssp.* | indica | 109 | -1.403 |
| *indica ssp.* | indica | 110 | -0.490 |
| *indica ssp.* | indica | 117 | -1.000 |
| *indica ssp.* | indica | 119 | -0.173 |
| *indica ssp.* | indica | 123 | -0.170 |
| *indica ssp.* | indica | 125 | -0.780 |
| *indica ssp.* | indica | 126 | -0.947 |
| *indica ssp.* | indica | 129 | -0.723 |
| *indica ssp.* | indica | 130 | -0.255 |
| *indica ssp.* | indica | 132 | -0.177 |
| *indica ssp.* | indica | 136 | 2.210 |
| *indica ssp.* | indica | 138 | -1.287 |
| *indica ssp.* | indica | 142 | 1.060 |
| *indica ssp.* | indica | 145 | -1.937 |
| *indica ssp.* | indica | 146 | -0.620 |
| *indica ssp.* | indica | 148 | -1.130 |
| *indica ssp.* | indica | 156 | -0.720 |
| *indica ssp.* | indica | 159 | -1.073 |
| *indica ssp.* | indica | 161 | -0.533 |
| *indica ssp.* | indica | 162 | -0.237 |
| *indica ssp.* | indica | 163 | -0.623 |
| *indica ssp.* | indica | 171 | -0.470 |
| *indica ssp.* | indica | 172 | -0.795 |
| *indica ssp.* | indica | 196 | -0.900 |
| *indica ssp.* | indica | 203 | 0.583 |
| *indica ssp.* | indica | 207 | 0.243 |
| *indica ssp.* | indica | 208 | -1.183 |
| *indica ssp.* | indica | 209 | -0.017 |
| *indica ssp.* | indica | 222 | -1.580 |
| *indica ssp.* | indica | 231 | -0.425 |
| *indica ssp.* | indica | 234 | -0.810 |
| *indica ssp.* | indica | 235 | -0.643 |
| *indica ssp.* | indica | 241 | -0.190 |
| *indica ssp.* | indica | 252 | -1.163 |
| *indica ssp.* | indica | 254 | -0.273 |
| *indica ssp.* | indica | 255 | -0.207 |
| *indica ssp.* | indica | 269 | -0.993 |
| *indica ssp.* | indica | 284 | -1.143 |
| *indica ssp.* | indica | 298 | 0.123 |
| *indica ssp.* | indica | 299 | -0.360 |
| *indica ssp.* | indica | 304 | -1.523 |
| *indica ssp.* | indica | 313 | -0.997 |
| *indica ssp.* | indica | 315 | -0.233 |
| *indica ssp.* | indica | 325 | -1.770 |
| *indica ssp.* | indica | 337 | 0.195 |
| *indica ssp.* | indica | 339 | -0.107 |

| | | | |
|---|---|---|---|
| *indica ssp.* | indica | 348 | -0.760 |
| *indica ssp.* | indica | 349 | 0.513 |
| *indica ssp.* | indica | 356 | -1.467 |
| *indica ssp.* | indica | 385 | -0.520 |
| *indica ssp.* | indica | 616 | -0.350 |
| *indica ssp.* | indica | 620 | -1.357 |
| *indica ssp.* | indica | 623 | -1.087 |
| *indica ssp.* | indica | 626 | -1.690 |
| *indica ssp.* | indica | 633 | -0.190 |
| *indica ssp.* | indica | 634 | 0.240 |
| *indica ssp.* | indica | 636 | -1.627 |
| *indica ssp.* | indica | 642 | -0.890 |
| *indica ssp.* | indica | 643 | -1.940 |
| *indica ssp.* | indica | 644 | 1.440 |
| *indica ssp.* | indica | 648 | -0.813 |
| *japonica ssp.* | temperate-japonica | 279 | 0.400 |
| *japonica ssp.* | temperate-japonica | 121 | 0.683 |
| *japonica ssp.* | temperate-japonica | 265 | 0.400 |
| *japonica ssp.* | temperate-japonica | 281 | -0.665 |
| *japonica ssp.* | temperate-japonica | 1 | 0.977 |
| *japonica ssp.* | temperate-japonica | 9 | 1.725 |
| *japonica ssp.* | temperate-japonica | 10 | 0.763 |
| *japonica ssp.* | temperate-japonica | 15 | 0.135 |
| *japonica ssp.* | temperate-japonica | 31 | 2.510 |
| *japonica ssp.* | temperate-japonica | 32 | 0.073 |
| *japonica ssp.* | temperate-japonica | 36 | -0.220 |
| *japonica ssp.* | temperate-japonica | 51 | 0.130 |
| *japonica ssp.* | temperate-japonica | 52 | -0.220 |
| *japonica ssp.* | temperate-japonica | 56 | 1.560 |
| *japonica ssp.* | temperate-japonica | 62 | -0.060 |
| *japonica ssp.* | temperate-japonica | 63 | 1.407 |
| *japonica ssp.* | temperate-japonica | 64 | 1.460 |
| *japonica ssp.* | temperate-japonica | 67 | -0.233 |
| *japonica ssp.* | temperate-japonica | 79 | 1.837 |
| *japonica ssp.* | temperate-japonica | 83 | 1.060 |
| *japonica ssp.* | temperate-japonica | 86 | 0.283 |
| *japonica ssp.* | temperate-japonica | 91 | 0.547 |
| *japonica ssp.* | temperate-japonica | 94 | 0.230 |
| *japonica ssp.* | temperate-japonica | 103 | 1.300 |
| *japonica ssp.* | temperate-japonica | 104 | 0.030 |
| *japonica ssp.* | temperate-japonica | 113 | 1.507 |
| *japonica ssp.* | temperate-japonica | 115 | 0.493 |
| *japonica ssp.* | temperate-japonica | 118 | 2.677 |
| *japonica ssp.* | temperate-japonica | 133 | -0.833 |
| *japonica ssp.* | temperate-japonica | 134 | 0.183 |
| *japonica ssp.* | temperate-japonica | 144 | 1.667 |
| *japonica ssp.* | temperate-japonica | 151 | 1.847 |

| | | | |
|---|---|---|---|
| *japonica ssp.* | temperate-japonica | 154 | 1.757 |
| *japonica ssp.* | temperate-japonica | 155 | 0.477 |
| *japonica ssp.* | temperate-japonica | 157 | 0.053 |
| *japonica ssp.* | temperate-japonica | 158 | -0.870 |
| *japonica ssp.* | temperate-japonica | 169 | -0.695 |
| *japonica ssp.* | temperate-japonica | 173 | 0.777 |
| *japonica ssp.* | temperate-japonica | 177 | 0.567 |
| *japonica ssp.* | temperate-japonica | 179 | 3.447 |
| *japonica ssp.* | temperate-japonica | 180 | -0.073 |
| *japonica ssp.* | temperate-japonica | 181 | 0.400 |
| *japonica ssp.* | temperate-japonica | 184 | 0.837 |
| *japonica ssp.* | temperate-japonica | 186 | 0.827 |
| *japonica ssp.* | temperate-japonica | 192 | 1.300 |
| *japonica ssp.* | temperate-japonica | 204 | 0.417 |
| *japonica ssp.* | temperate-japonica | 216 | 0.010 |
| *japonica ssp.* | temperate-japonica | 219 | -0.397 |
| *japonica ssp.* | temperate-japonica | 220 | -0.233 |
| *japonica ssp.* | temperate-japonica | 224 | 0.397 |
| *japonica ssp.* | temperate-japonica | 225 | -0.077 |
| *japonica ssp.* | temperate-japonica | 232 | 0.300 |
| *japonica ssp.* | temperate-japonica | 233 | 0.227 |
| *japonica ssp.* | temperate-japonica | 243 | 0.153 |
| *japonica ssp.* | temperate-japonica | 245 | -0.657 |
| *japonica ssp.* | temperate-japonica | 247 | 0.923 |
| *japonica ssp.* | temperate-japonica | 248 | -0.557 |
| *japonica ssp.* | temperate-japonica | 250 | 0.507 |
| *japonica ssp.* | temperate-japonica | 256 | 0.377 |
| *japonica ssp.* | temperate-japonica | 257 | 0.113 |
| *japonica ssp.* | temperate-japonica | 263 | 0.480 |
| *japonica ssp.* | temperate-japonica | 267 | -0.020 |
| *japonica ssp.* | temperate-japonica | 268 | -0.315 |
| *japonica ssp.* | temperate-japonica | 275 | 0.215 |
| *japonica ssp.* | temperate-japonica | 277 | -0.093 |
| *japonica ssp.* | temperate-japonica | 282 | 0.357 |
| *japonica ssp.* | temperate-japonica | 283 | 0.763 |
| *japonica ssp.* | temperate-japonica | 287 | 1.203 |
| *japonica ssp.* | temperate-japonica | 288 | 0.257 |
| *japonica ssp.* | temperate-japonica | 289 | -0.305 |
| *japonica ssp.* | temperate-japonica | 290 | 0.903 |
| *japonica ssp.* | temperate-japonica | 291 | -0.227 |
| *japonica ssp.* | temperate-japonica | 292 | 0.350 |
| *japonica ssp.* | temperate-japonica | 295 | -0.153 |
| *japonica ssp.* | temperate-japonica | 296 | 0.340 |
| *japonica ssp.* | temperate-japonica | 297 | 1.590 |
| *japonica ssp.* | temperate-japonica | 300 | 1.215 |
| *japonica ssp.* | temperate-japonica | 301 | 0.773 |
| *japonica ssp.* | temperate-japonica | 302 | 0.180 |

| | | | |
|---|---|---|---|
| *japonica ssp.* | temperate-japonica | 303 | -0.303 |
| *japonica ssp.* | temperate-japonica | 306 | 0.980 |
| *japonica ssp.* | temperate-japonica | 307 | -0.460 |
| *japonica ssp.* | temperate-japonica | 311 | 1.943 |
| *japonica ssp.* | temperate-japonica | 333 | -0.703 |
| *japonica ssp.* | temperate-japonica | 334 | -0.190 |
| *japonica ssp.* | temperate-japonica | 338 | 0.150 |
| *japonica ssp.* | temperate-japonica | 355 | 0.240 |
| *japonica ssp.* | temperate-japonica | 363 | 0.287 |
| *japonica ssp.* | temperate-japonica | 365 | 1.400 |
| *japonica ssp.* | temperate-japonica | 366 | 0.955 |
| *japonica ssp.* | temperate-japonica | 368 | 0.983 |
| *japonica ssp.* | temperate-japonica | 380 | 0.253 |
| *japonica ssp.* | temperate-japonica | 639 | 0.073 |
| *japonica ssp.* | tropical-japonica | 7 | 0.620 |
| *japonica ssp.* | tropical-japonica | 8 | -2.770 |
| *japonica ssp.* | tropical-japonica | 14 | -1.163 |
| *japonica ssp.* | tropical-japonica | 22 | 0.455 |
| *japonica ssp.* | tropical-japonica | 23 | 0.010 |
| *japonica ssp.* | tropical-japonica | 24 | -0.435 |
| *japonica ssp.* | tropical-japonica | 25 | -0.447 |
| *japonica ssp.* | tropical-japonica | 26 | -0.983 |
| *japonica ssp.* | tropical-japonica | 27 | 1.243 |
| *japonica ssp.* | tropical-japonica | 37 | -0.583 |
| *japonica ssp.* | tropical-japonica | 46 | -0.267 |
| *japonica ssp.* | tropical-japonica | 54 | -0.580 |
| *japonica ssp.* | tropical-japonica | 59 | -0.815 |
| *japonica ssp.* | tropical-japonica | 65 | -0.867 |
| *japonica ssp.* | tropical-japonica | 69 | 0.337 |
| *japonica ssp.* | tropical-japonica | 70 | 0.033 |
| *japonica ssp.* | tropical-japonica | 73 | 0.200 |
| *japonica ssp.* | tropical-japonica | 75 | -0.613 |
| *japonica ssp.* | tropical-japonica | 84 | -1.470 |
| *japonica ssp.* | tropical-japonica | 89 | -0.810 |
| *japonica ssp.* | tropical-japonica | 92 | -0.460 |
| *japonica ssp.* | tropical-japonica | 98 | -1.317 |
| *japonica ssp.* | tropical-japonica | 99 | -1.090 |
| *japonica ssp.* | tropical-japonica | 101 | 0.567 |
| *japonica ssp.* | tropical-japonica | 107 | -0.750 |
| *japonica ssp.* | tropical-japonica | 108 | -0.270 |
| *japonica ssp.* | tropical-japonica | 116 | -0.540 |
| *japonica ssp.* | tropical-japonica | 120 | -1.555 |
| *japonica ssp.* | tropical-japonica | 122 | -1.180 |
| *japonica ssp.* | tropical-japonica | 135 | -0.690 |
| *japonica ssp.* | tropical-japonica | 139 | -1.170 |
| *japonica ssp.* | tropical-japonica | 147 | -0.660 |
| *japonica ssp.* | tropical-japonica | 149 | -0.397 |

| | | | |
|---|---|---|---|
| *japonica ssp.* | tropical-japonica | 150 | -0.153 |
| *japonica ssp.* | tropical-japonica | 164 | 0.120 |
| *japonica ssp.* | tropical-japonica | 165 | 0.393 |
| *japonica ssp.* | tropical-japonica | 167 | -0.430 |
| *japonica ssp.* | tropical-japonica | 174 | -0.383 |
| *japonica ssp.* | tropical-japonica | 176 | -0.080 |
| *japonica ssp.* | tropical-japonica | 183 | -0.615 |
| *japonica ssp.* | tropical-japonica | 185 | -0.475 |
| *japonica ssp.* | tropical-japonica | 187 | -0.163 |
| *japonica ssp.* | tropical-japonica | 190 | -0.920 |
| *japonica ssp.* | tropical-japonica | 193 | 0.410 |
| *japonica ssp.* | tropical-japonica | 195 | 1.097 |
| *japonica ssp.* | tropical-japonica | 198 | -0.050 |
| *japonica ssp.* | tropical-japonica | 199 | 0.193 |
| *japonica ssp.* | tropical-japonica | 201 | -0.420 |
| *japonica ssp.* | tropical-japonica | 202 | -0.170 |
| *japonica ssp.* | tropical-japonica | 213 | 0.510 |
| *japonica ssp.* | tropical-japonica | 214 | 0.700 |
| *japonica ssp.* | tropical-japonica | 215 | 0.897 |
| *japonica ssp.* | tropical-japonica | 223 | 1.370 |
| *japonica ssp.* | tropical-japonica | 226 | 0.113 |
| *japonica ssp.* | tropical-japonica | 229 | 1.537 |
| *japonica ssp.* | tropical-japonica | 239 | -0.663 |
| *japonica ssp.* | tropical-japonica | 240 | 0.370 |
| *japonica ssp.* | tropical-japonica | 242 | -0.877 |
| *japonica ssp.* | tropical-japonica | 251 | 0.740 |
| *japonica ssp.* | tropical-japonica | 258 | -0.415 |
| *japonica ssp.* | tropical-japonica | 274 | 0.535 |
| *japonica ssp.* | tropical-japonica | 285 | 0.187 |
| *japonica ssp.* | tropical-japonica | 286 | 0.040 |
| *japonica ssp.* | tropical-japonica | 308 | -0.087 |
| *japonica ssp.* | tropical-japonica | 309 | -0.530 |
| *japonica ssp.* | tropical-japonica | 310 | 0.510 |
| *japonica ssp.* | tropical-japonica | 342 | -0.040 |
| *japonica ssp.* | tropical-japonica | 347 | -0.047 |
| *japonica ssp.* | tropical-japonica | 350 | -0.970 |
| *japonica ssp.* | tropical-japonica | 352 | -0.780 |
| *japonica ssp.* | tropical-japonica | 375 | -0.793 |
| *japonica ssp.* | tropical-japonica | 377 | -1.713 |
| *japonica ssp.* | tropical-japonica | 379 | 0.023 |
| *japonica ssp.* | tropical-japonica | 381 | -1.033 |
| *japonica ssp.* | tropical-japonica | 384 | -0.160 |
| *japonica ssp.* | tropical-japonica | 391 | -0.763 |
| *japonica ssp.* | tropical-japonica | 392 | -1.067 |
| *japonica ssp.* | tropical-japonica | 394 | 0.527 |
| *japonica ssp.* | tropical-japonica | 395 | -1.137 |
| *japonica ssp.* | tropical-japonica | 396 | -1.050 |

| | | | |
|---|---|---|---|
| *japonica ssp.* | tropical-japonica | 397 | -0.607 |
| *japonica ssp.* | tropical-japonica | 619 | 0.177 |
| *japonica ssp.* | tropical-japonica | 621 | -0.780 |
| *japonica ssp.* | tropical-japonica | 624 | -0.540 |
| *japonica ssp.* | tropical-japonica | 625 | -0.880 |
| *japonica ssp.* | tropical-japonica | 628 | 0.425 |
| *japonica ssp.* | tropical-japonica | 630 | -1.023 |
| *japonica ssp.* | tropical-japonica | 635 | -1.300 |
| *japonica ssp.* | tropical-japonica | 638 | -0.110 |
| *nn* | admixed | 41 | 1.217 |
| *nn* | admixed | 68 | -0.777 |
| *nn* | admixed | 80 | -0.400 |
| *nn* | admixed | 168 | -0.713 |
| *nn* | admixed | 272 | 0.273 |
| *nn* | admixed | 278 | -1.470 |
| *nn* | admixed | 340 | 0.230 |
| *nn* | admixed | 344 | 0.493 |
| *nn* | admixed | 390 | 0.333 |
| *indica ssp.* | admixed | 39 | 0.440 |
| *indica ssp.* | admixed | 294 | -0.413 |
| *japonica ssp.* | admixed | 205 | 2.005 |
| *japonica ssp.* | admixed | 249 | 0.923 |
| *japonica ssp.* | admixed | 270 | 0.553 |
| *japonica ssp.* | admixed | 271 | -0.717 |
| *japonica ssp.* | admixed | 343 | 0.090 |
| *japonica ssp.* | admixed | 96 | -0.510 |
| *japonica ssp.* | admixed | 140 | -0.195 |
| *japonica ssp.* | admixed | 273 | -0.733 |
| *japonica ssp.* | admixed | 280 | -0.553 |
| *japonica ssp.* | admixed | 358 | 0.377 |
| *japonica ssp.* | admixed | 386 | 0.210 |
| *japonica ssp.* | admixed | 389 | 0.253 |
| *nn* | admixed | 197 | 0.407 |
| *indica ssp.* | admixed | 166 | 0.307 |
| *indica ssp.* | admixed | 206 | -1.363 |
| *indica ssp.* | admixed | 227 | -0.730 |
| *indica ssp.* | admixed | 259 | -0.757 |
| *indica ssp.* | admixed | 293 | -0.120 |
| *indica ssp.* | admixed | 332 | -1.520 |
| *japonica ssp.* | admixed | 20 | -0.230 |
| *japonica ssp.* | admixed | 40 | -0.490 |
| *japonica ssp.* | admixed | 55 | 0.663 |
| *japonica ssp.* | admixed | 60 | 0.080 |
| *japonica ssp.* | admixed | 87 | 0.567 |
| *japonica ssp.* | admixed | 100 | -0.290 |
| *japonica ssp.* | admixed | 114 | -0.423 |
| *japonica ssp.* | admixed | 128 | -0.117 |

| | | | |
|---|---|---|---|
| *japonica ssp.* | admixed | 182 | -0.020 |
| *japonica ssp.* | admixed | 211 | 0.003 |
| *japonica ssp.* | admixed | 217 | 0.700 |
| *japonica ssp.* | admixed | 218 | -0.263 |
| *japonica ssp.* | admixed | 236 | -0.283 |
| *japonica ssp.* | admixed | 237 | 0.037 |
| *japonica ssp.* | admixed | 244 | 0.093 |
| *japonica ssp.* | admixed | 253 | 0.160 |
| *japonica ssp.* | admixed | 264 | -0.883 |
| *japonica ssp.* | admixed | 266 | -0.363 |
| *japonica ssp.* | admixed | 305 | -0.457 |
| *japonica ssp.* | admixed | 335 | 0.860 |
| *japonica ssp.* | admixed | 364 | 1.215 |
| *japonica ssp.* | admixed | 376 | -1.687 |
| *japonica ssp.* | admixed | 387 | 0.970 |
| *japonica ssp.* | admixed | 388 | -1.033 |
| *japonica ssp.* | admixed | 618 | 0.787 |
| *japonica ssp.* | admixed | 622 | 1.137 |
| *japonica ssp.* | admixed | 627 | -0.413 |
| *japonica ssp.* | admixed | 629 | 0.847 |
| *nn* | unclassified | 95 | 0.570 |
| *nn* | unclassified | 617 | -0.515 |
| *japonica ssp.* | unclassified | 631 | 0.050 |
| *indica ssp.* | unclassified | 28 | 0.180 |
| *indica ssp.* | unclassified | 354 | -0.420 |
| *indica ssp.* | unclassified | 612 | 0.690 |
| *nn* | unclassified | 210 | -0.447 |
| *nn* | unclassified | 393 | -1.127 |
| *nn* | unclassified | 650 | -0.730 |
| *nn* | unclassified | 653 | -1.450 |
| *japonica ssp.* | unclassified | 2 | 2.060 |
| *japonica ssp.* | unclassified | 238 | -0.403 |
| *japonica ssp.* | unclassified | 770 | -0.327 |
