## Supplementary material for "Rice plants alter their yield component parameters based on their tolerance to low temperatures determined by a non-invasive method during the vegetative stage": TABLE S4

**TABLE S4** Predictive panicle weight ratio for RPD1 genotypes according their particular canonical discriminant scores

| subspecies | subpopulation | NSFTV ID# | PPW |
|---|---|---|---|
| *japonica ssp.* | aromatic | 5 | 0.934 |
| *japonica ssp.* | aromatic | 12 | 0.938 |
| *japonica ssp.* | aromatic | 16 | 0.951 |
| *japonica ssp.* | aromatic | 45 | 1.209 |
| *japonica ssp.* | aromatic | 53 | 1.041 |
| *japonica ssp.* | aromatic | 93 | 1.121 |
| *japonica ssp.* | aromatic | 112 | 0.934 |
| *japonica ssp.* | aromatic | 124 | 1.255 |
| *japonica ssp.* | aromatic | 160 | 1.055 |
| *japonica ssp.* | aromatic | 191 | 0.957 |
| *japonica ssp.* | aromatic | 221 | 1.196 |
| *japonica ssp.* | aromatic | 260 | 1.134 |
| *japonica ssp.* | aromatic | 373 | 1.102 |
| *japonica ssp.* | aromatic | 640 | 1.050 |
| *indica ssp.* | aus | 4 | 0.818 |
| *indica ssp.* | aus | 6 | 0.645 |
| *indica ssp.* | aus | 13 | 0.568 |
| *indica ssp.* | aus | 18 | 0.706 |
| *indica ssp.* | aus | 19 | 0.447 |
| *indica ssp.* | aus | 33 | 0.653 |
| *indica ssp.* | aus | 44 | 0.880 |
| *indica ssp.* | aus | 49 | 0.865 |
| *indica ssp.* | aus | 50 | 0.683 |
| *indica ssp.* | aus | 58 | 1.288 |
| *indica ssp.* | aus | 78 | 0.962 |
| *indica ssp.* | aus | 81 | 0.897 |
| *indica ssp.* | aus | 85 | 0.577 |
| *indica ssp.* | aus | 88 | 0.658 |
| *indica ssp.* | aus | 105 | 0.853 |
| *indica ssp.* | aus | 131 | 1.285 |
| *indica ssp.* | aus | 152 | 0.896 |
| *indica ssp.* | aus | 153 | 0.789 |
| *indica ssp.* | aus | 178 | 0.885 |
| *indica ssp.* | aus | 200 | 0.781 |
| *indica ssp.* | aus | 228 | 0.963 |
| *indica ssp.* | aus | 246 | 1.163 |
| *indica ssp.* | aus | 261 | 0.627 |
| *indica ssp.* | aus | 262 | 0.948 |
| *indica ssp.* | aus | 276 | 0.893 |
| *indica ssp.* | aus | 312 | 0.796 |
| *indica ssp.* | aus | 314 | 0.916 |
| *indica ssp.* | aus | 316 | 0.849 |
| *indica ssp.* | aus | 317 | 0.641 |

| | | | |
|---|---|---|---|
| *indica ssp.* | aus | 318 | 0.847 |
| *indica ssp.* | aus | 319 | 0.873 |
| *indica ssp.* | aus | 320 | 0.870 |
| *indica ssp.* | aus | 321 | 0.657 |
| *indica ssp.* | aus | 322 | 0.939 |
| *indica ssp.* | aus | 323 | 0.652 |
| *indica ssp.* | aus | 324 | 0.699 |
| *indica ssp.* | aus | 326 | 0.874 |
| *indica ssp.* | aus | 327 | 0.813 |
| *indica ssp.* | aus | 328 | 0.836 |
| *indica ssp.* | aus | 329 | 0.901 |
| *indica ssp.* | aus | 330 | 0.728 |
| *indica ssp.* | aus | 331 | 0.874 |
| *indica ssp.* | aus | 336 | 0.669 |
| *indica ssp.* | aus | 341 | 0.704 |
| *indica ssp.* | aus | 345 | 0.981 |
| *indica ssp.* | aus | 346 | 0.908 |
| *indica ssp.* | aus | 353 | 0.722 |
| *indica ssp.* | aus | 357 | 0.903 |
| *indica ssp.* | aus | 359 | 0.787 |
| *indica ssp.* | aus | 360 | 0.939 |
| *indica ssp.* | aus | 369 | 0.793 |
| *indica ssp.* | aus | 370 | 0.787 |
| *indica ssp.* | aus | 371 | 0.746 |
| *indica ssp.* | aus | 372 | 0.738 |
| *indica ssp.* | aus | 378 | 0.805 |
| *indica ssp.* | aus | 637 | 0.511 |
| *indica ssp.* | indica | 3 | 0.861 |
| *indica ssp.* | indica | 11 | 0.849 |
| *indica ssp.* | indica | 17 | 0.565 |
| *indica ssp.* | indica | 21 | 0.879 |
| *indica ssp.* | indica | 29 | 0.779 |
| *indica ssp.* | indica | 30 | 0.931 |
| *indica ssp.* | indica | 34 | 0.743 |
| *indica ssp.* | indica | 35 | 0.815 |
| *indica ssp.* | indica | 43 | 0.946 |
| *indica ssp.* | indica | 57 | 0.904 |
| *indica ssp.* | indica | 61 | 0.883 |
| *indica ssp.* | indica | 66 | 1.022 |
| *indica ssp.* | indica | 71 | 0.887 |
| *indica ssp.* | indica | 72 | 0.880 |
| *indica ssp.* | indica | 74 | 0.638 |
| *indica ssp.* | indica | 76 | 0.671 |
| *indica ssp.* | indica | 77 | 0.584 |
| *indica ssp.* | indica | 90 | 0.914 |
| *indica ssp.* | indica | 97 | 0.673 |
| *indica ssp.* | indica | 102 | 0.733 |

| | | | |
|---|---|---|---|
| *indica ssp.* | indica | 106 | 0.534 |
| *indica ssp.* | indica | 109 | 0.564 |
| *indica ssp.* | indica | 110 | 0.802 |
| *indica ssp.* | indica | 117 | 0.669 |
| *indica ssp.* | indica | 119 | 0.885 |
| *indica ssp.* | indica | 123 | 0.886 |
| *indica ssp.* | indica | 125 | 0.726 |
| *indica ssp.* | indica | 126 | 0.683 |
| *indica ssp.* | indica | 129 | 0.741 |
| *indica ssp.* | indica | 130 | 0.863 |
| *indica ssp.* | indica | 132 | 0.884 |
| *indica ssp.* | indica | 136 | 1.507 |
| *indica ssp.* | indica | 138 | 0.594 |
| *indica ssp.* | indica | 142 | 1.207 |
| *indica ssp.* | indica | 145 | 0.424 |
| *indica ssp.* | indica | 146 | 0.768 |
| *indica ssp.* | indica | 148 | 0.635 |
| *indica ssp.* | indica | 156 | 0.742 |
| *indica ssp.* | indica | 159 | 0.650 |
| *indica ssp.* | indica | 161 | 0.791 |
| *indica ssp.* | indica | 162 | 0.868 |
| *indica ssp.* | indica | 163 | 0.767 |
| *indica ssp.* | indica | 171 | 0.807 |
| *indica ssp.* | indica | 172 | 0.722 |
| *indica ssp.* | indica | 196 | 0.695 |
| *indica ssp.* | indica | 203 | 1.082 |
| *indica ssp.* | indica | 207 | 0.994 |
| *indica ssp.* | indica | 208 | 3.224 |
| *indica ssp.* | indica | 209 | 0.926 |
| *indica ssp.* | indica | 222 | 0.517 |
| *indica ssp.* | indica | 231 | 0.819 |
| *indica ssp.* | indica | 234 | 0.719 |
| *indica ssp.* | indica | 235 | 0.762 |
| *indica ssp.* | indica | 241 | 0.880 |
| *indica ssp.* | indica | 252 | 0.626 |
| *indica ssp.* | indica | 254 | 0.859 |
| *indica ssp.* | indica | 255 | 0.876 |
| *indica ssp.* | indica | 269 | 0.671 |
| *indica ssp.* | indica | 284 | 0.631 |
| *indica ssp.* | indica | 298 | 0.962 |
| *indica ssp.* | indica | 299 | 0.836 |
| *indica ssp.* | indica | 304 | 0.532 |
| *indica ssp.* | indica | 313 | 0.670 |
| *indica ssp.* | indica | 315 | 0.869 |
| *indica ssp.* | indica | 325 | 0.468 |
| *indica ssp.* | indica | 337 | 0.981 |
| *indica ssp.* | indica | 339 | 0.902 |

| | | | |
|---|---|---|---|
| *indica ssp.* | indica | 348 | 0.732 |
| *indica ssp.* | indica | 349 | 1.064 |
| *indica ssp.* | indica | 356 | 0.547 |
| *indica ssp.* | indica | 385 | 0.794 |
| *indica ssp.* | indica | 616 | 0.839 |
| *indica ssp.* | indica | 620 | 0.576 |
| *indica ssp.* | indica | 623 | 0.646 |
| *indica ssp.* | indica | 626 | 0.489 |
| *indica ssp.* | indica | 633 | 0.880 |
| *indica ssp.* | indica | 634 | 0.993 |
| *indica ssp.* | indica | 636 | 0.505 |
| *indica ssp.* | indica | 642 | 0.698 |
| *indica ssp.* | indica | 643 | 0.423 |
| *indica ssp.* | indica | 644 | 1.306 |
| *indica ssp.* | indica | 648 | 0.718 |
| *japonica ssp.* | temperate-japonica | 279 | 1.035 |
| *japonica ssp.* | temperate-japonica | 121 | 1.109 |
| *japonica ssp.* | temperate-japonica | 265 | 1.035 |
| *japonica ssp.* | temperate-japonica | 281 | 0.756 |
| *japonica ssp.* | temperate-japonica | 1 | 1.185 |
| *japonica ssp.* | temperate-japonica | 9 | 1.381 |
| *japonica ssp.* | temperate-japonica | 10 | 1.129 |
| *japonica ssp.* | temperate-japonica | 15 | 0.965 |
| *japonica ssp.* | temperate-japonica | 31 | 1.586 |
| *japonica ssp.* | temperate-japonica | 32 | 0.949 |
| *japonica ssp.* | temperate-japonica | 36 | 0.873 |
| *japonica ssp.* | temperate-japonica | 51 | 0.964 |
| *japonica ssp.* | temperate-japonica | 52 | 0.873 |
| *japonica ssp.* | temperate-japonica | 56 | 1.338 |
| *japonica ssp.* | temperate-japonica | 62 | 0.914 |
| *japonica ssp.* | temperate-japonica | 63 | 1.298 |
| *japonica ssp.* | temperate-japonica | 64 | 1.311 |
| *japonica ssp.* | temperate-japonica | 67 | 0.869 |
| *japonica ssp.* | temperate-japonica | 79 | 1.410 |
| *japonica ssp.* | temperate-japonica | 83 | 1.207 |
| *japonica ssp.* | temperate-japonica | 86 | 1.004 |
| *japonica ssp.* | temperate-japonica | 91 | 1.073 |
| *japonica ssp.* | temperate-japonica | 94 | 0.990 |
| *japonica ssp.* | temperate-japonica | 103 | 1.270 |
| *japonica ssp.* | temperate-japonica | 104 | 0.938 |
| *japonica ssp.* | temperate-japonica | 113 | 1.324 |
| *japonica ssp.* | temperate-japonica | 115 | 1.059 |
| *japonica ssp.* | temperate-japonica | 118 | 1.629 |
| *japonica ssp.* | temperate-japonica | 133 | 0.712 |
| *japonica ssp.* | temperate-japonica | 134 | 0.978 |
| *japonica ssp.* | temperate-japonica | 144 | 1.365 |
| *japonica ssp.* | temperate-japonica | 151 | 1.412 |

| | | | |
|---|---|---|---|
| *japonica ssp.* | temperate-japonica | 154 | 1.389 |
| *japonica ssp.* | temperate-japonica | 155 | 1.055 |
| *japonica ssp.* | temperate-japonica | 157 | 0.944 |
| *japonica ssp.* | temperate-japonica | 158 | 0.703 |
| *japonica ssp.* | temperate-japonica | 169 | 0.749 |
| *japonica ssp.* | temperate-japonica | 173 | 1.133 |
| *japonica ssp.* | temperate-japonica | 177 | 1.078 |
| *japonica ssp.* | temperate-japonica | 179 | 1.830 |
| *japonica ssp.* | temperate-japonica | 180 | 0.911 |
| *japonica ssp.* | temperate-japonica | 181 | 1.035 |
| *japonica ssp.* | temperate-japonica | 184 | 1.149 |
| *japonica ssp.* | temperate-japonica | 186 | 1.146 |
| *japonica ssp.* | temperate-japonica | 192 | 1.270 |
| *japonica ssp.* | temperate-japonica | 204 | 1.039 |
| *japonica ssp.* | temperate-japonica | 216 | 1.125 |
| *japonica ssp.* | temperate-japonica | 219 | 0.826 |
| *japonica ssp.* | temperate-japonica | 220 | 0.869 |
| *japonica ssp.* | temperate-japonica | 224 | 1.034 |
| *japonica ssp.* | temperate-japonica | 225 | 0.910 |
| *japonica ssp.* | temperate-japonica | 232 | 1.008 |
| *japonica ssp.* | temperate-japonica | 233 | 1.114 |
| *japonica ssp.* | temperate-japonica | 243 | 0.970 |
| *japonica ssp.* | temperate-japonica | 245 | 0.759 |
| *japonica ssp.* | temperate-japonica | 247 | 1.171 |
| *japonica ssp.* | temperate-japonica | 248 | 0.785 |
| *japonica ssp.* | temperate-japonica | 250 | 1.062 |
| *japonica ssp.* | temperate-japonica | 256 | 1.028 |
| *japonica ssp.* | temperate-japonica | 257 | 0.960 |
| *japonica ssp.* | temperate-japonica | 263 | 1.055 |
| *japonica ssp.* | temperate-japonica | 267 | 0.925 |
| *japonica ssp.* | temperate-japonica | 268 | 0.848 |
| *japonica ssp.* | temperate-japonica | 275 | 0.986 |
| *japonica ssp.* | temperate-japonica | 277 | 0.906 |
| *japonica ssp.* | temperate-japonica | 282 | 1.023 |
| *japonica ssp.* | temperate-japonica | 283 | 1.129 |
| *japonica ssp.* | temperate-japonica | 287 | 1.244 |
| *japonica ssp.* | temperate-japonica | 288 | 0.997 |
| *japonica ssp.* | temperate-japonica | 289 | 0.850 |
| *japonica ssp.* | temperate-japonica | 290 | 1.166 |
| *japonica ssp.* | temperate-japonica | 291 | 0.871 |
| *japonica ssp.* | temperate-japonica | 292 | 1.022 |
| *japonica ssp.* | temperate-japonica | 295 | 0.890 |
| *japonica ssp.* | temperate-japonica | 296 | 1.019 |
| *japonica ssp.* | temperate-japonica | 297 | 1.345 |
| *japonica ssp.* | temperate-japonica | 300 | 1.247 |
| *japonica ssp.* | temperate-japonica | 301 | 1.132 |
| *japonica ssp.* | temperate-japonica | 302 | 0.977 |

| | | | |
|---|---|---|---|
| *japonica ssp.* | temperate-japonica | 303 | 0.851 |
| *japonica ssp.* | temperate-japonica | 306 | 1.186 |
| *japonica ssp.* | temperate-japonica | 307 | 0.810 |
| *japonica ssp.* | temperate-japonica | 311 | 1.438 |
| *japonica ssp.* | temperate-japonica | 333 | 0.746 |
| *japonica ssp.* | temperate-japonica | 334 | 0.880 |
| *japonica ssp.* | temperate-japonica | 338 | 0.969 |
| *japonica ssp.* | temperate-japonica | 355 | 0.993 |
| *japonica ssp.* | temperate-japonica | 363 | 1.005 |
| *japonica ssp.* | temperate-japonica | 365 | 1.296 |
| *japonica ssp.* | temperate-japonica | 366 | 1.180 |
| *japonica ssp.* | temperate-japonica | 368 | 1.187 |
| *japonica ssp.* | temperate-japonica | 380 | 0.996 |
| *japonica ssp.* | temperate-japonica | 639 | 0.949 |
| *japonica ssp.* | tropical-japonica | 7 | 1.092 |
| *japonica ssp.* | tropical-japonica | 8 | 0.207 |
| *japonica ssp.* | tropical-japonica | 14 | 0.626 |
| *japonica ssp.* | tropical-japonica | 22 | 1.049 |
| *japonica ssp.* | tropical-japonica | 23 | 0.809 |
| *japonica ssp.* | tropical-japonica | 24 | 0.816 |
| *japonica ssp.* | tropical-japonica | 25 | 0.813 |
| *japonica ssp.* | tropical-japonica | 26 | 0.673 |
| *japonica ssp.* | tropical-japonica | 27 | 1.255 |
| *japonica ssp.* | tropical-japonica | 37 | 0.778 |
| *japonica ssp.* | tropical-japonica | 46 | 0.860 |
| *japonica ssp.* | tropical-japonica | 54 | 0.779 |
| *japonica ssp.* | tropical-japonica | 59 | 0.717 |
| *japonica ssp.* | tropical-japonica | 65 | 0.704 |
| *japonica ssp.* | tropical-japonica | 69 | 1.018 |
| *japonica ssp.* | tropical-japonica | 70 | 0.939 |
| *japonica ssp.* | tropical-japonica | 73 | 0.982 |
| *japonica ssp.* | tropical-japonica | 75 | 0.770 |
| *japonica ssp.* | tropical-japonica | 84 | 0.546 |
| *japonica ssp.* | tropical-japonica | 89 | 0.719 |
| *japonica ssp.* | tropical-japonica | 92 | 0.810 |
| *japonica ssp.* | tropical-japonica | 98 | 0.586 |
| *japonica ssp.* | tropical-japonica | 99 | 0.645 |
| *japonica ssp.* | tropical-japonica | 101 | 1.078 |
| *japonica ssp.* | tropical-japonica | 107 | 0.734 |
| *japonica ssp.* | tropical-japonica | 108 | 0.860 |
| *japonica ssp.* | tropical-japonica | 116 | 0.789 |
| *japonica ssp.* | tropical-japonica | 120 | 0.524 |
| *japonica ssp.* | tropical-japonica | 122 | 0.622 |
| *japonica ssp.* | tropical-japonica | 135 | 0.750 |
| *japonica ssp.* | tropical-japonica | 139 | 0.624 |
| *japonica ssp.* | tropical-japonica | 147 | 0.758 |
| *japonica ssp.* | tropical-japonica | 149 | 0.826 |

| | | | |
|---|---|---|---|
| *japonica ssp.* | tropical-japonica | 150 | 0.890 |
| *japonica ssp.* | tropical-japonica | 164 | 0.741 |
| *japonica ssp.* | tropical-japonica | 165 | 1.033 |
| *japonica ssp.* | tropical-japonica | 167 | 0.818 |
| *japonica ssp.* | tropical-japonica | 174 | 0.830 |
| *japonica ssp.* | tropical-japonica | 176 | 0.909 |
| *japonica ssp.* | tropical-japonica | 183 | 0.769 |
| *japonica ssp.* | tropical-japonica | 185 | 0.806 |
| *japonica ssp.* | tropical-japonica | 187 | 0.887 |
| *japonica ssp.* | tropical-japonica | 190 | 0.690 |
| *japonica ssp.* | tropical-japonica | 193 | 1.037 |
| *japonica ssp.* | tropical-japonica | 195 | 1.217 |
| *japonica ssp.* | tropical-japonica | 198 | 0.917 |
| *japonica ssp.* | tropical-japonica | 199 | 0.981 |
| *japonica ssp.* | tropical-japonica | 201 | 0.820 |
| *japonica ssp.* | tropical-japonica | 202 | 0.886 |
| *japonica ssp.* | tropical-japonica | 213 | 1.063 |
| *japonica ssp.* | tropical-japonica | 214 | 1.113 |
| *japonica ssp.* | tropical-japonica | 215 | 1.164 |
| *japonica ssp.* | tropical-japonica | 223 | 1.288 |
| *japonica ssp.* | tropical-japonica | 226 | 0.960 |
| *japonica ssp.* | tropical-japonica | 229 | 1.331 |
| *japonica ssp.* | tropical-japonica | 239 | 0.757 |
| *japonica ssp.* | tropical-japonica | 240 | 1.027 |
| *japonica ssp.* | tropical-japonica | 242 | 0.701 |
| *japonica ssp.* | tropical-japonica | 251 | 1.123 |
| *japonica ssp.* | tropical-japonica | 258 | 0.822 |
| *japonica ssp.* | tropical-japonica | 274 | 1.070 |
| *japonica ssp.* | tropical-japonica | 285 | 0.979 |
| *japonica ssp.* | tropical-japonica | 286 | 0.941 |
| *japonica ssp.* | tropical-japonica | 308 | 0.907 |
| *japonica ssp.* | tropical-japonica | 309 | 0.792 |
| *japonica ssp.* | tropical-japonica | 310 | 1.063 |
| *japonica ssp.* | tropical-japonica | 342 | 0.920 |
| *japonica ssp.* | tropical-japonica | 347 | 0.918 |
| *japonica ssp.* | tropical-japonica | 350 | 0.677 |
| *japonica ssp.* | tropical-japonica | 352 | 0.726 |
| *japonica ssp.* | tropical-japonica | 375 | 0.723 |
| *japonica ssp.* | tropical-japonica | 377 | 0.483 |
| *japonica ssp.* | tropical-japonica | 379 | 0.936 |
| *japonica ssp.* | tropical-japonica | 381 | 0.660 |
| *japonica ssp.* | tropical-japonica | 384 | 0.888 |
| *japonica ssp.* | tropical-japonica | 391 | 0.731 |
| *japonica ssp.* | tropical-japonica | 392 | 0.651 |
| *japonica ssp.* | tropical-japonica | 394 | 1.068 |
| *japonica ssp.* | tropical-japonica | 395 | 0.633 |
| *japonica ssp.* | tropical-japonica | 396 | 0.656 |

| | | | |
|---|---|---|---|
| *japonica ssp.* | tropical-japonica | 397 | 0.772 |
| *japonica ssp.* | tropical-japonica | 619 | 0.976 |
| *japonica ssp.* | tropical-japonica | 621 | 0.726 |
| *japonica ssp.* | tropical-japonica | 624 | 0.789 |
| *japonica ssp.* | tropical-japonica | 625 | 0.700 |
| *japonica ssp.* | tropical-japonica | 628 | 1.041 |
| *japonica ssp.* | tropical-japonica | 630 | 0.663 |
| *japonica ssp.* | tropical-japonica | 635 | 0.591 |
| *japonica ssp.* | tropical-japonica | 638 | 0.901 |
| *nn* | admixed | 41 | 1.248 |
| *nn* | admixed | 68 | 0.727 |
| *nn* | admixed | 80 | 0.826 |
| *nn* | admixed | 168 | 0.744 |
| *nn* | admixed | 272 | 1.001 |
| *nn* | admixed | 278 | 0.546 |
| *nn* | admixed | 340 | 0.990 |
| *nn* | admixed | 344 | 1.059 |
| *nn* | admixed | 390 | 1.017 |
| *indica ssp.* | admixed | 39 | 1.045 |
| *indica ssp.* | admixed | 294 | 0.822 |
| *japonica ssp.* | admixed | 205 | 1.454 |
| *japonica ssp.* | admixed | 249 | 1.171 |
| *japonica ssp.* | admixed | 270 | 1.075 |
| *japonica ssp.* | admixed | 271 | 0.743 |
| *japonica ssp.* | admixed | 343 | 0.954 |
| *japonica ssp.* | admixed | 96 | 0.797 |
| *japonica ssp.* | admixed | 140 | 0.879 |
| *japonica ssp.* | admixed | 273 | 0.739 |
| *japonica ssp.* | admixed | 280 | 0.786 |
| *japonica ssp.* | admixed | 358 | 1.028 |
| *japonica ssp.* | admixed | 386 | 0.985 |
| *japonica ssp.* | admixed | 389 | 0.996 |
| *nn* | admixed | 197 | 1.036 |
| *indica ssp.* | admixed | 166 | 1.010 |
| *indica ssp.* | admixed | 206 | 0.574 |
| *indica ssp.* | admixed | 227 | 0.739 |
| *indica ssp.* | admixed | 259 | 0.732 |
| *indica ssp.* | admixed | 293 | 0.899 |
| *indica ssp.* | admixed | 332 | 0.533 |
| *japonica ssp.* | admixed | 20 | 0.870 |
| *japonica ssp.* | admixed | 40 | 0.802 |
| *japonica ssp.* | admixed | 55 | 1.103 |
| *japonica ssp.* | admixed | 60 | 0.951 |
| *japonica ssp.* | admixed | 87 | 1.078 |
| *japonica ssp.* | admixed | 100 | 0.854 |
| *japonica ssp.* | admixed | 114 | 0.820 |
| *japonica ssp.* | admixed | 128 | 0.900 |

| | | | |
|---|---|---|---|
| *japonica ssp.* | admixed | 182 | 0.925 |
| *japonica ssp.* | admixed | 211 | 0.931 |
| *japonica ssp.* | admixed | 217 | 1.113 |
| *japonica ssp.* | admixed | 218 | 0.861 |
| *japonica ssp.* | admixed | 236 | 0.856 |
| *japonica ssp.* | admixed | 237 | 0.940 |
| *japonica ssp.* | admixed | 244 | 0.954 |
| *japonica ssp.* | admixed | 253 | 0.972 |
| *japonica ssp.* | admixed | 264 | 0.699 |
| *japonica ssp.* | admixed | 266 | 0.835 |
| *japonica ssp.* | admixed | 305 | 0.811 |
| *japonica ssp.* | admixed | 335 | 1.155 |
| *japonica ssp.* | admixed | 364 | 1.247 |
| *japonica ssp.* | admixed | 376 | 0.490 |
| *japonica ssp.* | admixed | 387 | 1.183 |
| *japonica ssp.* | admixed | 388 | 0.660 |
| *japonica ssp.* | admixed | 618 | 1.136 |
| *japonica ssp.* | admixed | 622 | 1.227 |
| *japonica ssp.* | admixed | 627 | 0.822 |
| *japonica ssp.* | admixed | 629 | 1.151 |
| *nn* | others | 95 | 1.079 |
| *nn* | others | 617 | 0.796 |
| *japonica ssp.* | others | 631 | 0.943 |
| *indica ssp.* | others | 28 | 0.977 |
| *indica ssp.* | others | 354 | 0.820 |
| *indica ssp.* | others | 612 | 1.110 |
| *nn* | others | 210 | 0.813 |
| *nn* | others | 393 | 0.636 |
| *nn* | others | 650 | 0.739 |
| *nn* | others | 653 | 0.551 |
| *japonica ssp.* | others | 2 | 1.468 |
| *japonica ssp.* | others | 238 | 0.825 |
| *japonica ssp.* | others | 770 | 0.845 |
